# SMARCA4 deficient tumours are vulnerable to KDM6A/UTX and KDM6B/JMJD3 blockade

**DOI:** 10.1101/2020.08.07.241232

**Authors:** Octavio A. Romero, Andrea Vilarrubi, Juan J. Alburquerque-Bejar, Antonio Gomez, Alvaro Andrades, Deborah Trastulli, Eva Pros, Sara Verdura, Lourdes Farré, Juan F. Martín-Tejera, Paula Llabata, Ana Oaknin, Maria Saigi, Josep M. Piulats, Xavier Matias-Guiu, Pedro P. Medina, August Vidal, Alberto Villanueva, Montse Sanchez-Cespedes

## Abstract

Despite the genetic inactivation of *SMARCA4*, a core component of the SWI/SNF-complex commonly found in cancer, there are no therapies that effectively target SMARCA4-deficient tumours. Here, we show that, unlike the cells with activated *MYC* oncogene, cells with *SMARCA4* inactivation are refractory to the histone deacetylase inhibitor, SAHA, leading to the aberrant accumulation of H3K27me3. This is associated with impaired transactivation and significantly reduced levels of the histone demethylases KDM6A/UTX and KDM6B/JMJD3, which confer a strong dependency on the KDM6s in the *SMARCA4*-mutant cells, so that its inhibition compromises cell viability. Administering the KDM6 inhibitor GSK-J4 to mice orthotopically implanted with *SMARCA4*-mutant lung cancer cells or primary small cell carcinoma of the hypercalcaemic type (SCCOHT) had strong anti-tumour effects. Our results highlight the vulnerability of KDM6 inhibitors as a characteristic that could be exploited for treating *SMARCA4-*mutant cancer patients.

## Introduction

Chromatin remodelling is one of the epigenetic processes that is commonly disturbed in cancer, mainly through alterations in the mammalian switch/sucrose non-fermentable (SWI/SNF) complex. This complex modifies the structure of the chromatin by the ATP-dependent disruption of DNA–histone interactions at the nucleosomes, thereby activating or repressing gene expression. The various functions and components of the SWI/SNF complex have been thoroughly reviewed elsewhere^1,2^.

Alterations at genes encoding different components of the SWI/SNF complex are present in a variety of tumour types and are thus an important feature of cancer development^3^. The *SMARCA4* (also known as *BRG1*) gene codes a core catalytic component of the SWI/SNF complex that features a bromodomain and helicase/ATPase activity^1,2^. Our own previous work produced the first evidence that *SMARCA4* is genetically inactivated in cancer and that SMARCA4 deficiency prevents the response to pro-differentiation stimuli in cancer cells^4-6^. In lung cancer, *SMARCA4* inactivation affects about one-third of non-small cell lung cancers (NSCLCs) and preferentially occurs against a background of wild type MYC (either C, L or N) or of members of the MYC-axis, such as MAX or MGA^4-7^. This hints at the existence of an important network that connects SWI/SNF and MAX/MYC functions. Mutations of *SMARCA4* also occur in other types of cancer, notably in the rare and very aggressive small cell carcinoma of the ovary, hypercalcaemic type (SCCOHT)^8^, in which *SMARCA4* inactivation has been reported in almost 100% of cases^9-11^.

The progress made towards understanding the role of chromatin remodelling in cancer development highlights the great potential of new epigenetic-based therapeutic strategies. With particular reference to SMARCA4, some previous studies have sought the vulnerabilities of SMARCA4-deficient tumours with a view to exploiting them for cancer treatment. SMARCA4 and SMARCA2 are mutually exclusive catalytic subunits of the SWI/SNF complex, and the inhibition of SMARCA2 activity appears to be synthetic lethal in cancer cells carrying *SMARCA4*-inactivating mutations, an effect that could be explained by paralogue insufficiency^12-13^. Further, SWI/SNF-mutant cancer cells with a wild type *KRAS* background depend on the non-catalytic action of the histone methyltransferase, EZH2^14^. However, we currently know of no small compounds that are capable of suppressing the ATPase or non-catalytic functions of SMARCA2 and EZH2, respectively, so these molecules are not yet suitable for use in therapeutic interventions. More recently, it has been proposed that cancer cells with an inactive SMARCA4 may be susceptible to CDK4/6 inhibitors^15^.

On the other hand, components of the SWI/SNF complex bind to various nuclear receptors (e.g., oestrogen, androgen, glucocorticoid and retinoid receptors), thereby adapting the gene expression programs to the demands of the cell environment^16-19^. We have reported that SMARCA4 is required to promote cell growth inhibition triggered by corticoids and retinoids in cancer cells^6^, and that such effects are enhanced by combination with the pan-histone deacetylase (HDAC) inhibitor suberanilohydroxamic acid (SAHA)^20^. We observed that *MYC*-amplified but not *SMARCA4*-mutant cancer cells, were sensitive to these treatments.

Here, we found that defective regulation of gene expression causes cancer cells that lack SMARCA4 to exhibit very low levels of KDM6s. This forms not only the basis of the refractoriness to SAHA, but also sensitizes the cancer cells to inhibition of the demethylase activity of KDM6s, heavily compromising their viability.

## Results

### Refractoriness to growth inhibition and increase in H3K27me3 by SAHA in SMARCA4-deficient cells

We studied the differential response to SAHA in lung cancer cells with oncogenic activation of MYC, in other words, with respect to the high levels of expression due to gene amplification (hereafter referred to as MYCamp) and genetic inactivation of SMARCA4 (hereafter, SMARCA4def). The administration of SAHA was more effective at reducing the growth of MYCamp cells than of SMARCA4def cells (mean of half-maximum effective concentrations (EC_50_) of 0.5 µM and 1.4 µM, for each group, respectively) (**Fig. 1a-b, Extended Data Fig. 1a**). These effects were not influenced by lung cancer histopathological subtypes, since they occurred in non-small (NSCLC) and small (SCLC) cell lung cancer types. The selective sensitivity to SAHA of MYC-oncogenic cells was validated using publicly available datasets including more than 750 cancer cell lines of different origin and genetic background (**Fig. 1c)**.

**Fig. 1.**
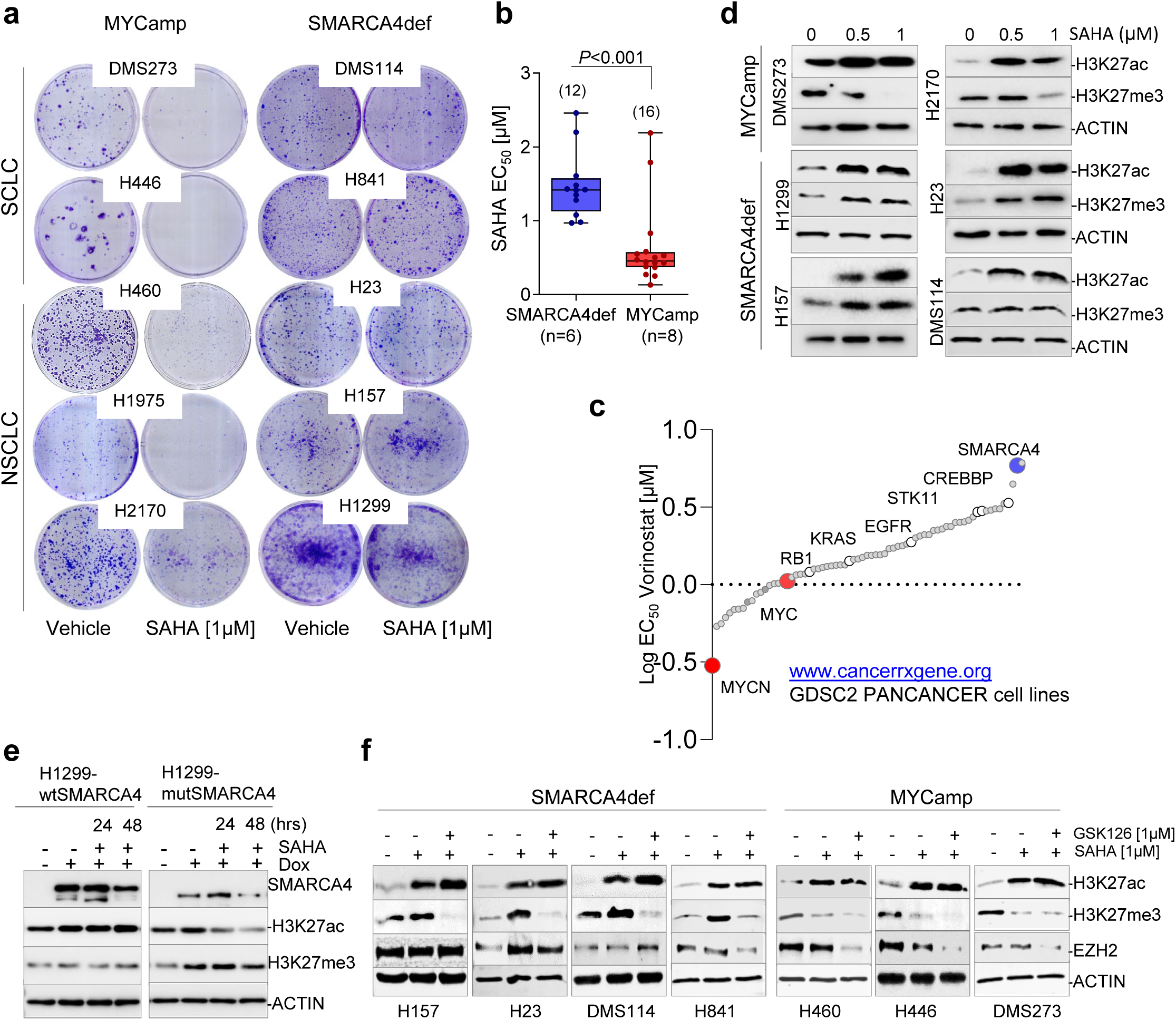
SAHA reduces growth and H3K27me3 levels in MYCamp, but not in SMARCA4def cancer cells. **a**, Clonogenic assays of the indicated lung cancer cells, untreated (vehicle) or treated with SAHA. **b**, Distribution and mean of EC_50_ values for treatment with SAHA in MYCamp (0.5 µM) and SMARCA4def cells (1.4 µM) (from MTT assays for cell viability; Extended Data Fig. 1). Bars show mean ± SD; two-sided unpaired Student’s t-test. Values from two independent experiments per cell line are represented. **c**, Plot with the comparative EC_50_ values for treatment with SAHA (vorinostat) in a panel of 758 cancer cell lines (from www.cancerrxgene.org database), according to the presence of selected gene alterations, including amplification of *MYC* oncogenes and inactivation of *SMARCA4*. **d, e, f**, Representative western blots depicting global levels of the indicated proteins and histone marks (H3K27ac and H3K27me3) in the indicated cells following treatment with SAHA (**d**), in the H1299-wtSMARCA4 and H1299-mutSMARCA4 cell models after induction of SMARCA4 (doxycycline, 1 µg/mL; 72 h), with or without SAHA treatment at 24 and 48 h (**e**), or in the indicated cells following treatments with SAHA and SAHA plus GSK126 (**f**). ACTIN, protein-loading control.

SAHA increases global histone acetylation, favouring an open chromatin structure and promoting transcriptional activation^21^. Transcriptionally active chromatin domains are characterized by a distinct array of histone marks, e.g., H3K27ac, H3K4me1 and H3K4me3, whereas H3K27me3 and H2AK119ub are often found at silent gene loci^22,23^. We first tested the effects of SAHA in global H3K27ac and H3K27me3, two different marks on the same residue, but with opposite functions^22-23^. As expected, SAHA triggered an increase in global H3K27ac in all the cells, while we observed an aberrant accumulation of H3K27me3 in the SMARCA4def cells, rather than the expected decrease (**Fig. 1d**). To determine whether this was due to a defective SMARCA4, we used H1299 cells, which lack SMARCA4 expression owing to an intragenic homozygous deletion^4^, and restituted wild type SMARCA4 (H1299-wtSMARCA4), using a doxycycline-inducible system, as previously reported^6^. As a control, we expressed a mutant form that lacked the ATPase domain (p.Glu668_Gln758del)^4^ (hereafter referred to as H1299-mutSMARCA4). Administration of SAHA did not affect global H3K27me3 in the H1299-wtSMARCA4 cells, whereas the H1299-mutSMARCA4 cells underwent an increase in H3K27me3 concomitantly with a decrease in global H3K27ac **(Fig. 1e)**. Thus, the absence of a functional SMARCA4 induces defects in the dynamics of the H3K27me3 mark, following administration of SAHA.

The net levels of H3K27me3 are dictated by the coordinated action of histone methyltransferases (EZH2) and demethylases (KDM6A and KDM6B)^23^. The administration of SAHA did not alter the levels of EZH2 in most SMARCA4def cells, indicating that overactivation of the methyltransferase activity of EZH2 is unlikely to account for the defects in H3K27me3 triggered by SAHA in these cells **(Fig. 1f; Extended Data Fig. 1b)**. In the MYCamp cells, SAHA treatment alone reduced the EZH2 levels, the effect being enhanced by the addition of GSK126, an inhibitor of the enzymatic activity of EZH2. The administration of GSK126, alone or in combination with SAHA, did not reduce the proliferation or viability of any group of cells **(Extended Data Fig. 1c-e)**, implying that EZH2 activity does not by itself cause the refractoriness to cell growth inhibition by SAHA in SMARCA4def cells.

### Regulation of KDM6 expression by SMARCA4

The histone demethylases KDM6A (also known as UTX) and KDM6B (also known as JMJD3) have H3K27 as a substrate and play a central role in the development of some types of tumours^23^. Searching the Cancer Cell Line Encyclopedia (CCLE) we found significantly lower levels of expression of several histone demethylases (KDMs), including *KDM6A* and *KDM6B*, in SMARCA4def compared with MYCamp cells **(Fig**.**2a)** (**Extended Data Fig. 2a)**. We validated our observations in a panel of cell lines at the mRNA and protein levels **(Fig**.**2b-c)**. Consistent with the low levels of KDM6s, basal global H3K27ac was present at a lower level in the SMARCA4def cells than in the MYCamp cells. The levels of EZH2 were similar in the two groups **(Fig. 2b)**. Further, the ectopic expression of the SMARCA4 wild type (H1299-wtSMARCA4 cells), but not of the mutant (H1299-mutSMARCA4 cells), triggered the upregulation of *KDM6A* and *KDM6B* (**Fig. 2d; Extended Data Fig. 2b**). Conversely, in the MYCamp cells, the depletion of SMARCA4 reduced the levels of both KDM6s (**Extended Data Fig. 2c**). Collectively, these observations indicate that a functional SMARCA4 is required to activate KDM6 expression.

**Fig. 2.**
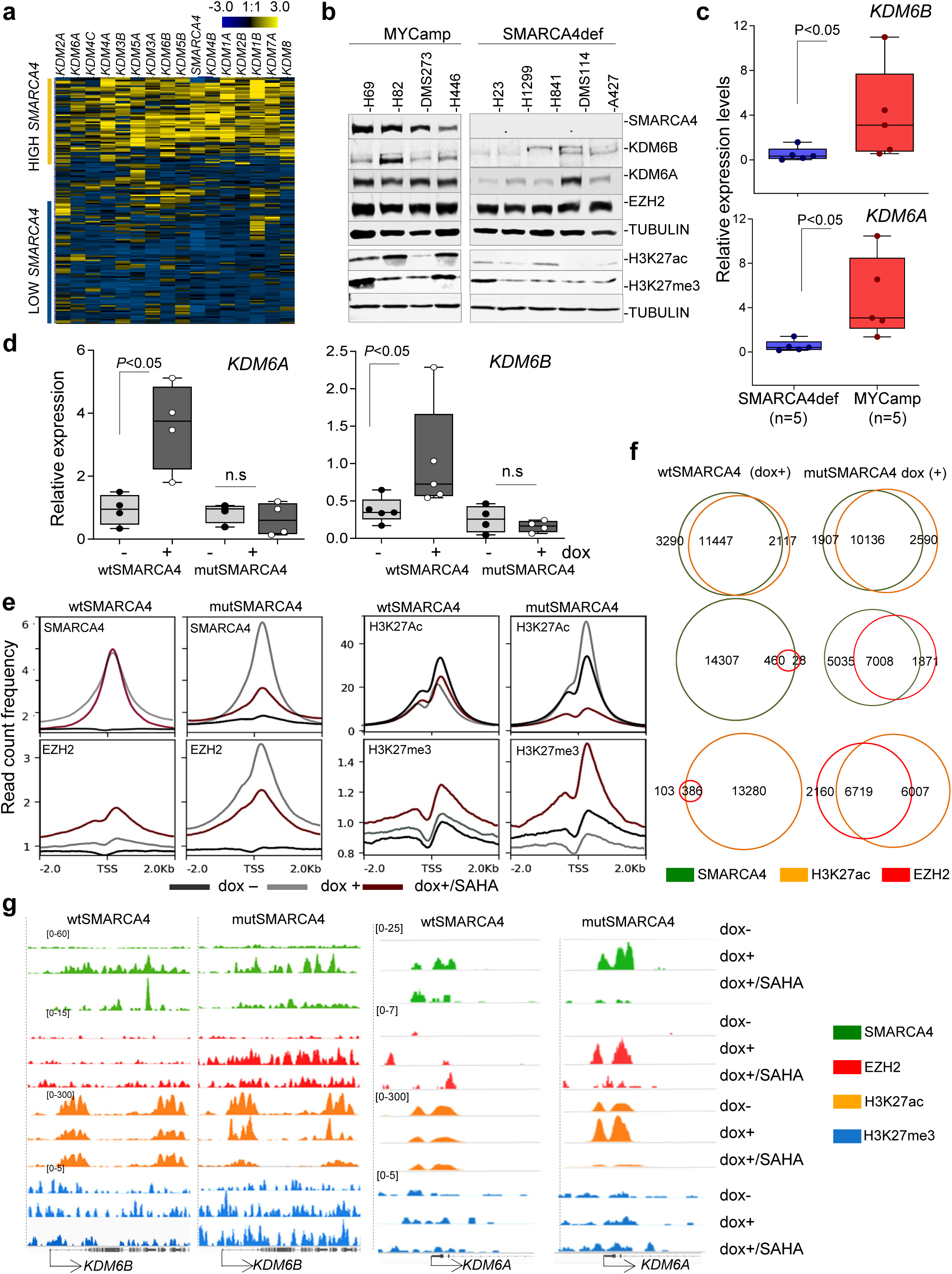
SMARCA4 regulates the levels of the KDM6s. **a**, Heatmap depicting mRNA levels (RPKMs, reads/kb/ million) of the indicated *KDMs* and of *SMARCA4* in 179 lung cancer cell lines (from Cancer Cell Line Encyclopedia-CCLE at cBioportal). **b**, Western blot depicting endogenous levels of the indicated proteins in lung cancer cell lines. TUBULIN, protein-loading control. **c, d**, Real-time quantitative PCR of *KDM6A* and *KDM6B* (relative to *ACTB*) for comparing mRNA levels among the indicated groups of lung cancer cell lines (**c**), or in the H1299-wtSMARCA4 and H1299-mutSMARCA4 cell models (doxycycline, 1 µg/mL; 72 h) (**d**). Bars show mean ± SD. Two-sided unpaired Student’s t-test, **e**, read count frequency of heatmaps, at ± 2 kb regions centred on the TSS, of the indicated proteins and conditions in the H1299 cell model (doxycycline, 1 µg/mL & SAHA 1 µM, for 72 h). **f**, Venn diagrams representing overlap of SMARCA4, H3K27ac and EZH2 peaks in the indicated cells and under the stipulated conditions. **g**, Representative snapshots from IGV of ChIP-seq profiles at selected target loci performed in H1299 cell models.

To study the genome-wide effects of wild type and mutant SMARCA4 and of SAHA on the dynamics of H3K27 modification, we performed chromatin immunoprecipitation followed by sequencing (ChIP-seq) of SMARCA4, EZH2, H3K27ac and H3K27me3, in the H1299 cell model. No peaks were observed for SMARCA4 before adding doxycycline, which is consistent with the absence of SMARCA4 in these cells (**Fig. 2e; Extended Data Fig. 3a**). The global occupancy of wild type and mutant SMARCA4 was similar, indicating that ATPase activity does not influence recruitment to the chromatin (**Fig. 2e-f; Extended Data Fig. 3a-b**). Also, H3K27ac deposition at promoters was not affected by SMARCA4 activity, despite there being a strong increase in EZH2 binding to the DNA in SMARCA4 mutant-expressing cells. The latter observation suggests that the overexpressed mutant protein has a dominant negative effect. H3K27ac marks were present in at least 80% of the promoters bound by SMARCA4 and some also showed EZH2 occupancy (**Fig. 2f; Extended Data Fig. 4a**). The increase in EZH2 binding following expression of the SMARCA4 mutant is consistent with previous findings that ectopic expression of an SMARCA4-inactive protein allows the occupancy of PRC1 and PRC2 in CpG island promoters throughout the genome^24^.

**Fig. 3.**
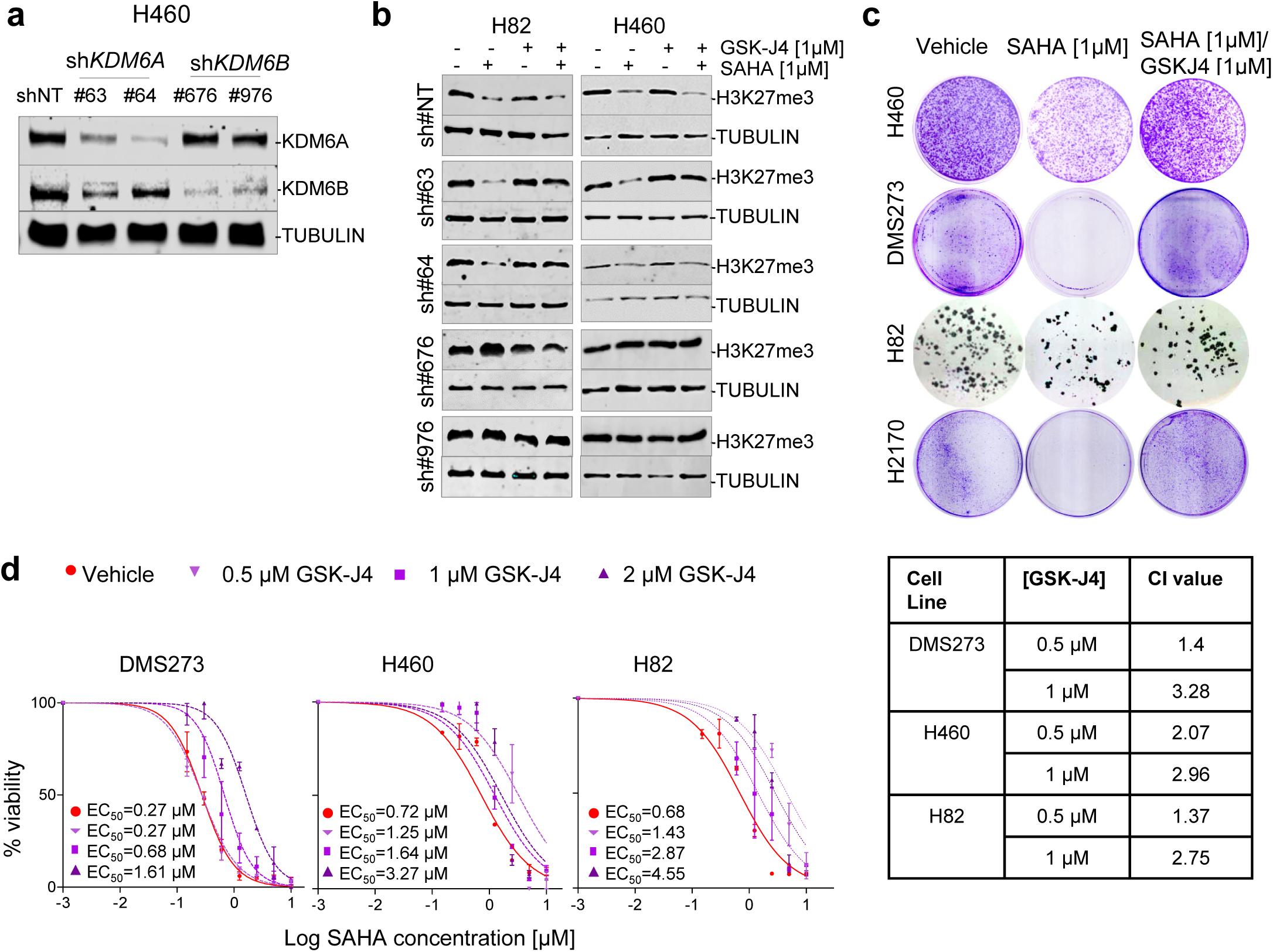
KDM6B depletion mimics the response of SMARCA4-deficient cells to SAHA. **a**, Western blot depicting levels of KDM6s in H460 cells infected with the non-target short hairpin (shNT) and with different shKDM6 (#63 and #64) and shKDM6B (#676, #976). **b**, Western blot depicting levels of H3K27me3 and cells infected with the shNT, shKDM6 (#63, #64) or shKDM6B (#676, #976) treated with GSK-J4 and/or SAHA for 72 h. **c**, Representative clonogenic assays for the indicated cells and treatments. **d**, Viability of indicated cell lines, measured using MTT assays, 5 days after treatment with increasing concentrations of SAHA and co-treated or not with GSK-J4 at different concentrations. Lines, number of viable cells relative to the total number at 0 h. Data are presented as mean ± SD from three replicate cell cultures in two experiments. On the right, table presenting the combination index (CI) at the indicated concentrations of GSK-J4 (average CI from two independent experiments). CI < 1, CI = 1 and CI > 1 indicate synergism, additive effect, and antagonism, respectively.

**Fig. 4.**
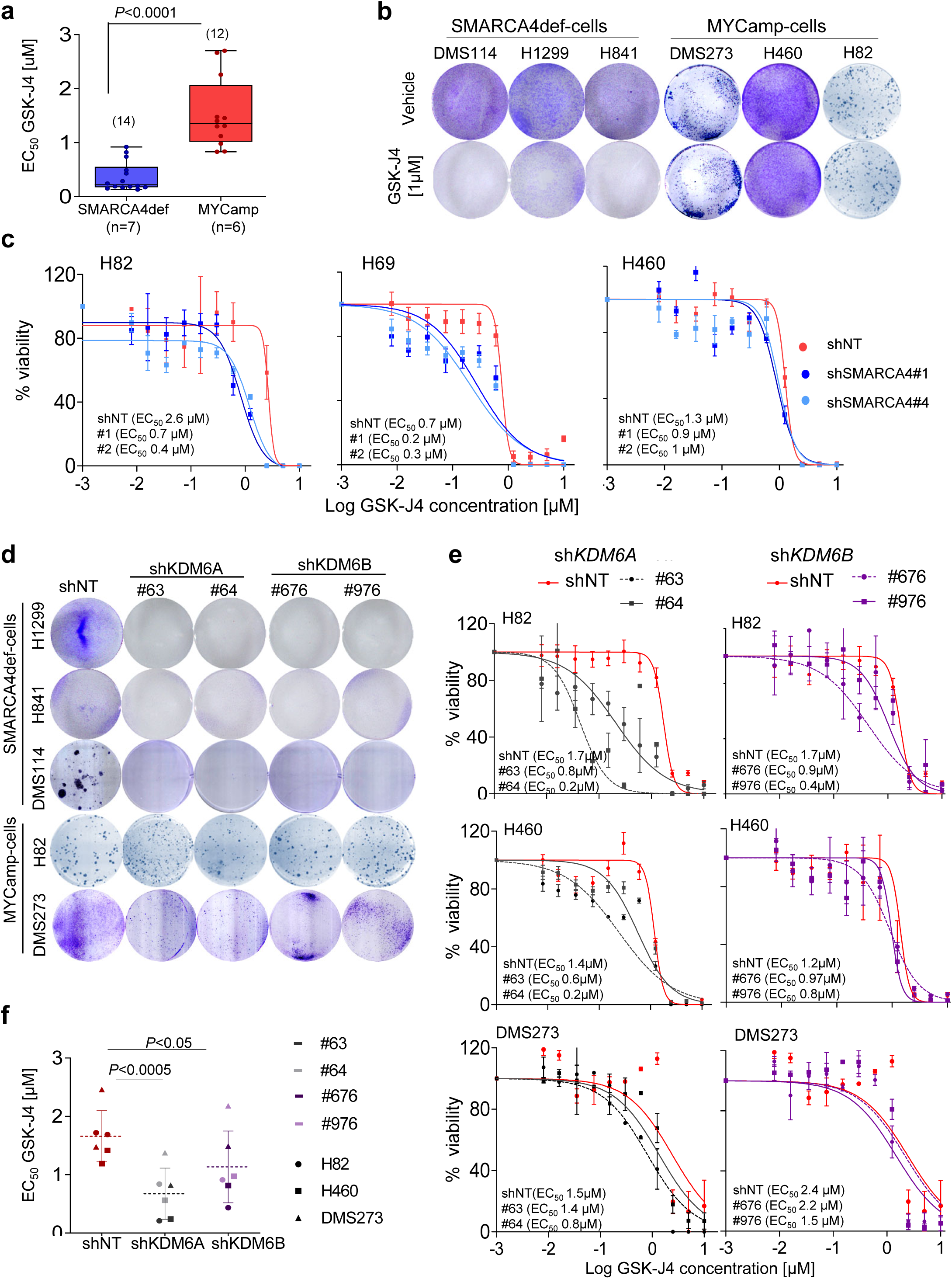
SMARCA4def cells are vulnerable to KDM6s inhibition. **a**, Distribution and mean of EC_50_ values for GSK-J4 (MTT assays. Extended Data Fig. 6) in the indicated groups of cells. Values, from each cell line and from two independent experiments are represented. Two-sided unpaired Student’s t-test. **b**, Representative clonogenic assays for the indicated cells and treatments. **c**, Viability of the indicated cells infected with a non-target (shNT) control or with two shRNAs targeting SMARCA4 (#1 & #4), measured using MTT assays, after treatment with increasing concentrations of GSK-J4 for 5 days. Lines, number of viable cells relative to the total number at 0 h. Data are presented as mean and SD from three replicates and two experiments. **d**, Representative clonogenic assays for the indicated cells infected with shNT, shKDM6 (#63, #64) or shKDM6B (#676, #976). **e**, Viability of the indicated cell lines, measured using MTT assays, infected with a non-target (shNT) control or with two shRNAs targeting KDM6A or KDM6B after treatment with increasing concentrations of GSK-J4 for 5 days. Lines, number of viable cells relative to the total number at 0 h. Error bars, mean ± SD from triplicates. **f**, Distribution and mean of the EC_50_ from two independent experiments. Lines show mean ± SD. *P*-values were calculated using paired two-tailed Student’s t test.

The administration of SAHA reduced the number of promoters recruiting SMARCA4 wild type by half, without producing major changes in global H3K27ac deposition (**Extended Data Fig. 3a-b)**. SMARCA4 does not interact directly with the DNA, but rather recognises and binds acetylated lysines within histone H3 and H4 tails^25^. Consistent with this, the severe reduction of global H3K27ac deposition observed in SMARCA4 mutant-expressing cells following SAHA treatment, also shown by western blot (**Fig. 1e**), was associated with a strong reduction in the global intensity of the SMARCA4 peaks (**Fig. 2e**). Conversely, the treatment with SAHA prompted the recruitment of EZH2 to the DNA and the increase in H3K27me3 deposition, in mutant and wild type SMARCA4-expressing cells. However, the effect in the latter cell type was less dramatic, and we attribute this to the slow rate of H3K27me3 removal (**Fig. 2e; Extended Data Fig. 3a**). The observation that SAHA increases EZH2 recruitment and H3K27me3 deposition is somewhat counterintuitive but may be a compensatory effect to avoid detrimentally high levels of histone acetylation.

SMARCA4 was bound to different KDMs, including *KDM2B, KDM4B, KDM6A* and *KDM6B*, among others, in association with H3K27ac. However, there was also a concomitant increase in EZH2 occupancy in the SMARCA4 mutant-expressing cells **(Fig. 2g; Extended Data Fig. 4b)**. These results support the idea that SMARCA4 regulates the expression of various KDMs, including *KDM6A* and *KDM6B*, through direct promoter occupancy. The increase in EZH2 in the promoter of these genes is consistent with a lack of transcriptional activation of these KDMs in the H1299-mutSMARCA4 cells **(Fig. 2b)**

### KDM6B depletion mimics the response of SMARCA4-deficient cells to SAHA

We wondered to what extent the lack of KDM6A or KDM6B regulation is involved in the greater H3K27me3 and refractoriness to SAHA in the SMARCA4def cells, and whether their relative contributions differ. First, we found that the mRNA levels of the *KDM6B* were inversely correlated with the EC_50_ to SAHA (**Extended Data Fig. 5a**). Next, using shRNAs, we downregulated KDM6A and KDM6B expression in different MYCamp cells **(Fig, 3a; Extended Fig. 5b)**, and noted that, mimicking the behaviour of the SMARCA4def cells, the reduction in *KDM6B* levels suppressed the ability to reduce the global level of H3K27me3 deposition **(Fig. 3b)** and to inhibit cell growth **(Extended Data Fig. 5c)** by SAHA. Conversely, the depletion of KDM6A did not affect these characteristics. The administration of the small molecule compound GSK-J4^26^, a very specific inhibitor of KDM6A/KDM6B reverted the sensitivity to SAHA **(Fig. 3c-d)**, and prevented the SAHA-triggered global decrease in H3K27me3 in the KDM6A-depleted cells (**Fig. 3b)**, in a dose-dependent manner.

**Fig. 5.**
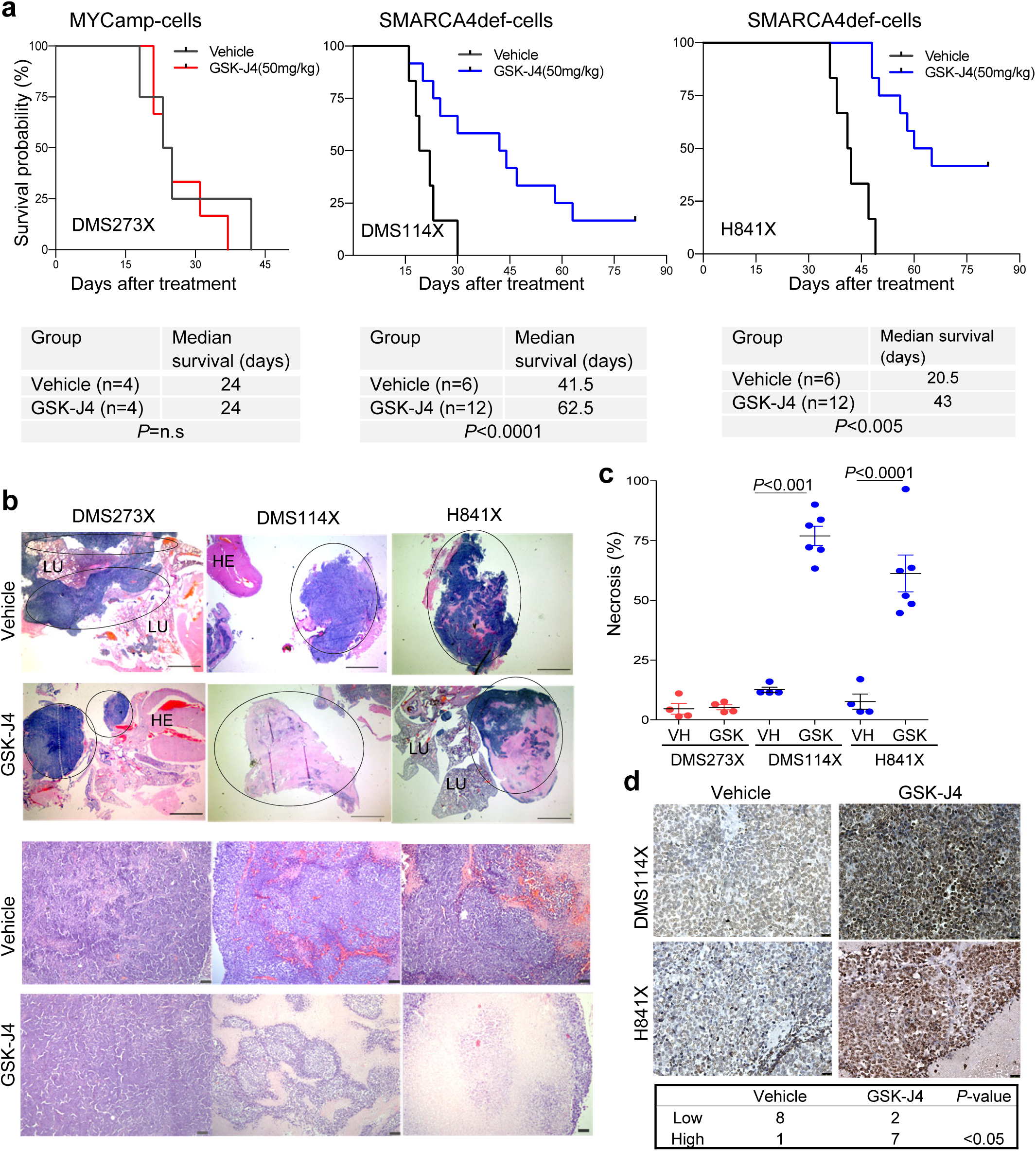
GSK-J4 induces tumour regression of SMARCA4def lung tumours *in vivo*. **a**, Kaplan-Meier curves showing overall survival for GSK-J4-treated compared with vehicle control groups of each indicated orthotopically implanted mice model. Panels below, number of mice (n) and mean survival times for each group of treatment and cell line. *P*-values from the two-sided log-rank (Mantel-Cox) test of the plots are included. n.s., not significant. **b**, Representative sections of haematoxylin and eosin (H&E) staining of tumours from the indicated cells, from mice treated with either GSK-4 or vehicle. Upper panels, tumour regions are marked within circles. Scale bars, 2.5 mm. The pink areas inside the tumours indicate necrosis. HE, heart; LU, lung. Bottom panels, representative sections, at higher magnification, of tumours from the indicated cells and treatments. Scale bars, 50 µm. **c**, Quantification of necrotic areas. Mean ± SD are indicated for each group. Two-sided unpaired Student’s t-test. VH, vehicle. **d**, Representative immunostaining of H3K27me3 in tumours from the indicated cells and treatments. Scale bars, 25 µm. Below, distribution of H3K27me3 staining among tumours (three tumours per cell line and condition) from the DMS273X, H841X and DMS114X tumours treated with vehicle or GSK-J4. Low (intensity values 1 & 2); high (intensity values 3 & 4). Fisher’s Exact test.

These findings suggest that the low levels of KDM6B account for the resistance of the SMARCA4def cells to growth inhibition by SAHA, and hint at a more widespread role for KDM6B in the global removal of H3K27me3.

### Inhibition of KDM6A/B is toxic in SMARCA4-deficient cancer cells

We hypothesized that the low levels and impaired regulation of KDM6s expression and the defects in H3K27 modification may render SMARCA4def cells particularly susceptible to KDM6s inhibition. We tested the effects of GSK-J4 on the growth of our panel of cancer cells. We found that the drug was more toxic in the SMARCA4def cells, with a five-fold lower EC_50_ than in the MYCamp cells **(Fig. 4a-b; Extended Data Fig. 6a)**. For the next stage of the study, we chose to use GSK-J4 at a concentration of 1 µM, which is the approximate mean of the EC_50_ of the MYCamp cells **(Fig. 4a)**. We depleted SMARCA4 in three MYCamp cells and observed a decrease in the EC_50_ for GSK-J4, which is further evidence that GSK-J4 is more toxic in cancer cells with a non-functional SMARCA4 (**Fig. 4c)**. We also tested the effects of rescuing SMARCA4 on the response to GSK-J4 using the H1299 cell model. Overexpression of the mutant SMARCA4 increased sensitivity to GSK-J4 relative to the restitution of wild type SMARCA4 (EC_50_, 0.11 µM *versus* 0.2 µM) (**Extended Data Fig. 6b-c)**. The toxicity was even greater in the H1299-mutSMARCA4 than in the parental H1299 cells, supporting the existence of a dominant negative effect of overexpressing a SMARCA4 mutant protein. GSK-J4 is a potent inhibitor of KDM6s but can also suppress the activity of other KDMs, so we investigated how the low levels of KDM6s suppress cell viability by depleting KDM6A- and KDM6B-. Downregulation of KDM6A or KDM6B inhibited the growth of the SMARCA4-def cells without affecting the MYCamp cells (**Fig. 4d)**. Moreover, the lower levels of KDM6A, and to a lesser extent of *KDM6B*, in the MYCamp cells sensitised the cells to the treatment with GSK-J4 (**Fig. 4e--f**). Together, these findings imply that the lack of SMARCA4 confers vulnerability to KDM6s inhibition on cancer cells, and that the intrinsically low levels of KDM6s, caused by the defective function of SMARCA4, underpin these effects.

**Fig. 6.**
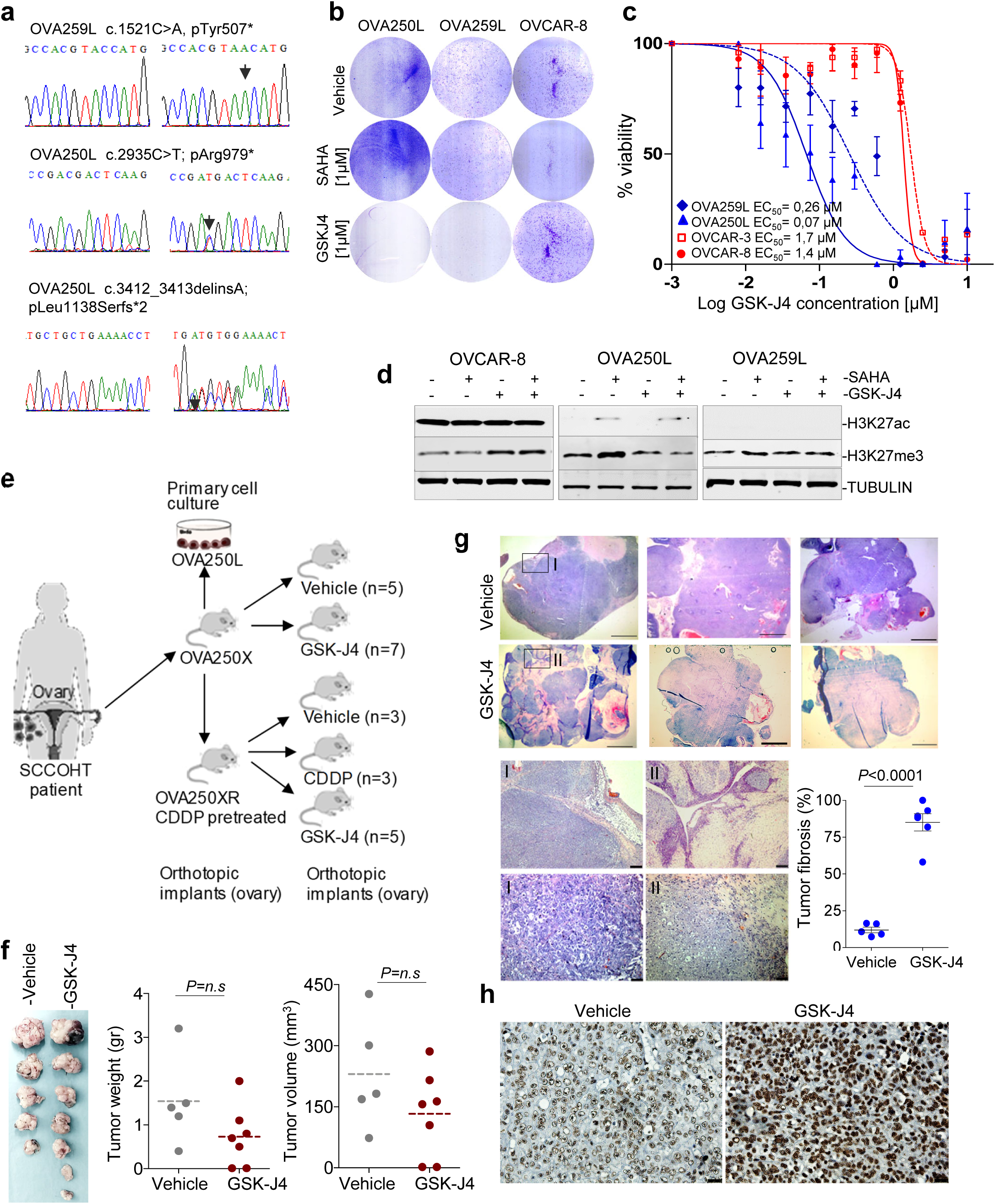
GSK-J4 reduces cancer cell viability of SCCOHT orthotopically implanted in mice. **a**, Chromatogram depicting changes in *SMARCA4*, at the genomic DNA level, in the SCCOHTs of two patients. The alterations were biallelic, confirming the complete inactivation of *SMARCA4*. A normal reference DNA is also included. **b**, Representative clonogenic assays for the indicated cells and treatments. **c**, Percentage of viable cells of the indicated cells, measured using MTT assays, after treatment with increasing concentrations of GSK-J4 for 5 days. Lines show the number of viable cells relative to the total number at 0 h. Error bars, mean ± SD from three replicates. **d**, Western blot of the endogenous levels of the indicated proteins and cancer cells. TUBULIN, protein-loading control. **e**, Schematic representation of different models and treatments in mice developed from the OVA-250 tumour. **f**, Left panel, gross pathological photographs at necropsy of the ovarian tumours that arose in mice treated with vehicle or GSK-J4. Right panels, volume and weight of each tumour. Two-sided unpaired Student’s t-test. n.s. not significant. **g**, Representative sections of H&E staining of tumours from the group of mice treated as indicated. Pink areas inside the tumours indicate fibrosis. Scale bars, 2.5 mm; Lower panels, magnification of the areas indicated (rectangle); scale bar, 50 µm (above) and 25 µm (below); Lower right panel, quantification of fibrotic areas. Two-sided unpaired Student’s t-test. **h**, Representative immunostaining of H3K27me3 for OVA250X tumours treated with vehicle or GSK-J4. Scale bars, 25 µm.

### Anti-tumour effects of GSK-J4 in *SMARCA4*def lung cancer orthotopic mouse models

We investigated the ability of the GSK-J4 compound to suppress tumour growth *in vivo*. To this end, we first grew two of the SMARCA4def (DMS114 and H841) and one MYCamp (DMS273) cell lines subcutaneously into the back of the mice (n = 3 mice/cell line). Once the solid tumour had entered the exponential growth phase, mice were euthanized, and the tumours we minced into small fragments and orthotopically implanted into the lungs of another cohort of mice, as previously described^20,27^, to generate the orthotopic tumours. We randomly assigned the animals, implanted with each of the tumours, to treatment or vehicle groups of mice. Treatment with GSK-J4 strongly increased the overall and median survival of the animals implanted with the SMARCA4def tumours (DMS114X and H841X) relative to their matched vehicle group, whereas we found no differences between vehicle and treated groups in the animals implanted with the MYCamp tumours (DMS273X) (**Fig. 5a**). Remarkably, five of the mice implanted with the H841X and two of those implanted with the DMS114X and treated with GSK-J4 were alive at the end of the experiment, but had to be sacrificed despite not having respiratory difficulties or other symptoms associated with tumour progression. Our histopathological examination of tumour masses revealed the existence of large areas of necrosis in the tumours from GSK-J4-treated mice in comparison with the tumours from the vehicle-treated mice (**Fig. 5b-c; Extended Data Fig. 7**). We also used immunohistochemistry to determine the changes in the levels of H3K27me3 in the tumour samples following treatment with GKS-J4. We noted a significant increase in the levels of H3K27me3 in all the tumours treated with GSK-J4, suggesting that the compound had effectively reached the tumours (**Fig. 5d**).

### GSK-J4 reduces tumour growth in mice implanted with SMARCA4-mutant SCCOHT

As previously mentioned, SCCOHT is a very aggressive and rare type of ovarian cancer that features inactivation of SMARCA4 in almost all cases^9-11^. Here, we generated patient derived orthotopic xenografts (PDOXs) using the primary tumours of two SCCOHT patients (OVA250 and OVA259), by orthotopically implanting the tumour in the mouse ovary, as previously described^28^. We used the PDOXs, in their first pass, to derive primary cancer cell cultures (OVA250L and OVA259L). We confirmed the presence of biallelic inactivating mutations at *SMARCA4* and the lack of protein in the two patients’ tumour cells and PDOXs **(Fig. 6a; Extended Data Fig. 8a)**. First, we tested the effects of the SAHA and GSK-J4 compounds in the primary cultures and included, as a reference, two commercial epithelial ovarian carcinoma cell lines (OVCAR-3 and OVCAR-8), which are wild type for *SMARCA4* (https://cancer.sanger.ac.uk/cell_lines). Treatment with GSK-J4 strongly suppressed cell viability and clonogenic capability, exclusively in the OVA250L and OVA259L cells, whereas SAHA did not affect cell growth (**Fig. 6b-c**). It was puzzling to observe that the OVA250L and OVA259L cells had extremely low levels of global H3K27ac, even after treatment with SAHA. Similar to what happened in the lung cancer cells, the global basal levels of H3K27me3 were increased after administration of SAHA **(Fig. 6d)**.

Next, we investigated the influence of GSK-J4 treatment on the growth of the OVA250 tumour *in vivo*. We orthotopically implanted primary tumours, either treatment-naïve or derived from mice previously treated with cisplatin (CDDP) (see Methods section for further details), in the ovary of female nude mice to generate the OVA250X and OVA250XR tumours, respectively **(Fig. 6e; Extended Data Fig. 8b-e)**. We observed a reduction in the size and weight of the tumours from mice treated with the GSK-J4 inhibitor in the OVA250X model **(Fig. 6f)**. Although the differences did not reach statistical significance, histological examination revealed the presence of a few viable tumour cells in the tumours from mice treated with GSK-J4. These tissues contained a large amount of fibrosis, instead of the necrosis observed in the lung cancer orthotopic model, possibly because the OVA250X was derived from a primary tumour and not from cancer cell lines **(Fig. 6g)**. Similar to what was observed in the lung cancer mouse models, the GSK-J4 treatment triggered an increase in H3K27me3 **(Fig. 6h)**. Next, we generated the OVA250XR tumours to determine the benefits of GSK-J4 in tumours that have been pre-treated with CDDP, the standard treatment for SCCOHT. The morphology of the OVA250XR tumours was similar with that of the primary tumour and of the OVA250X (**Extended Data Fig. 8d**). The OVA250XR showed greater refractoriness to CDDP than did OVA250X (**Extended Data Fig. 8e)**. The treatment with GSK-J4 significantly decreased tumour growth in the OVA250XR. Taken together, these observations demonstrate that the KDM6s inhibitor could constitute a therapeutic option for SCCOHT patients even after they have become resistant to CDDP (**Extended Data Fig. 8f-h)**.

## Discussion

Here, we present evidences that cancer cells carrying oncogenic MYC are susceptible to growth inhibition by treatment with the histone deacetylase (HDAC) inhibitor, SAHA. HDAC inhibitors, including SAHA, have come to be recognised as biologically active compounds of value for treating cancers, although their use is currently limited to some haematological malignancies^29^. Our current findings indicate that the pre-selection of patients with tumours in which any of the MYC family of genes have been genetically activated will have better response rates to SAHA, which suggests that SAHA could be used to treat neuroblastomas and lung cancers, among others types of cancer, in which the *MYC* genes are amplified.

Second, we found that lung or ovarian cancer cells with inactivated *SMARCA4* not only were refractory to the growth suppression triggered by SAHA, but also aberrantly accumulated H3K27me3 following the administration of this inhibitor. The levels of global H3K27ac were low in the SMARCA4def cells, a characteristic that was accentuated in the SCCOHT cells. We ruled out a central role for EZH2 methyltransferase activity in the refractoriness to SAHA in these cells, although the dependency on a non-catalytic role for EZH2 cannot be completely discounted, as it is known to affect the survival of the SWI/SNF-mutant cancer cells^14^. Additionally, we demonstrate that the transactivation of several lysine demethylases (KDMs), including KDM6A and KDM6B, is impaired in cells that lack SMARCA4, leading to a downregulation of basal KDM6s. This, coupled with inability of these cells to modulate the levels of EZH2 expression in response to SAHA and, in keeping with other knowledge^6,16-20^, is evidence that defective chromatin remodelling in SMARCA4def cells promotes a closed chromatin structure and a transcriptionally rigid scenario that maintains the refractoriness of these cells to the appropriate modification of gene expression upon different stimuli.

Despite the high degree of sequence similarity in the catalytic domain of KDM6A and KDM6B, these two enzymes have some specific roles^30^. KDM6A is mainly associated with the demethylation of H3K27me3 at the transcriptional start sites of the HOX genes upon differentiation stimuli, whereas KDM6B is involved in inflammation and other, general physiological processes^30^. Furthermore, KDM6A, but not KDM6B, is responsible for Kabuki syndrome (KS), an infrequent, inherited disease that is characterized by neurological, endocrine and autoimmune disorders^31^. Here, we found that low levels of KDM6B are responsible for the refractoriness to SAHA and the global increase in H3K27me3 upon administration of SAHA in SMARCA4def cells. This is consistent with a broader role for KMD6B in H3K27me3 deposition than that of KDM6A. In keeping with these observations, there are no known major global changes in H3K27me3, unlike in their wild type counterparts^32^. Non-catalytic activities have been proposed for the KDM6s which may also account for some of these differences^30^.

Considering its potential clinical applicability, the most relevant finding presented here is the great vulnerability of the SMARCA4def cells to KDM6s inhibition, which was evident in cell culture and in mouse models with orthotopic transplants of lung cancer cells and SCCOHTs. Their frequency and poor prognosis mean that the use of GSK-J4 or similar compounds can have a great impact on the treatment of SMARCA4-mutant tumours. Likewise, SCCOHT is an aggressive carcinoma with rhabdoid characteristics and, though infrequent overall, predominantly affects young women^8-11^. Our current results, which show that GSK-J4 strongly suppresses its growth *in vivo*, emphasises the huge potential of KDM6s inhibitors in treating this disease, which otherwise has very limited treatment options and, consequently, a dismal prognosis. Currently, there is only limited, preclinical information about the use of KDM6s inhibitors in cancer treatment. Anti-tumorigenic activities of GSK-J4 have been shown in some leukaemias and in gliomas with *H3F3A* mutations, both of which are attributable to the inhibition of KDM6B^33-34^. In our case, the depletion of either KDM6A or KDM6B affected the viability of SMARCA4def cells, suggesting that, in a cancer cell context with very low levels of KDM6s, the depletion of either of them cannot be compensated. We demonstrate that the intrinsically invariable low levels of KDM6s, which are due to the lack of SMARCA4-mediated regulation, are the underlying cause of the toxicity to the KDM6 inhibitor, GSK-J4, in SMARCA4def cells. Despite our current findings about the toxicity of the GSK-J4 compound in SMARCAdef cells, the effects could be broader, and cancer cells in which other members of the SWI/SNF-complex or of related pathways are genetically inactivated, may also be vulnerable to this inhibitor.

## Extended Data Figure Legends

**Extended Data Fig.1. a**, Representative examples of effects on cell viability, measured using MTT assays, in the indicated cell lines after treatment with increasing concentrations of SAHA for 5 days. **b**, Western blot depicting the levels of EZH2 in the indicated cancer cells, following SAHA treatment. ACTIN, protein-loading control. **c, d**, Effects on cell viability, measured using MTT assays, in the indicated cell lines after treatment with increasing concentrations of GSK-126 (**c**) or of SAHA and 1 µM of GSK-126, for 5 days (**d**). **e**, Representative clonogenic assays for the indicated cells and treatments.

**Extended Data Fig. 2. a**, Violin plots comparing levels of the indicated *KDMs* in SMARCA4def, MYCamp and other (wild type for *SMARCA4* and for *MYC*) cells, from a panel of 179 lung cancer cell lines (Cancer Cell Line Encyclopedia-CCLE at cBioportal). RPKMs (reads per million). Bars show mean ± SD. Two-sided unpaired Student’s t-test: **, *P* < 0.01; ***, *P* < 0.001; ****, *P* < 0.0001. **b**, Determination of protein levels of KDM6A and KDM6B by western blot in the H1299-wtSMARCA4 and H1299-mutSMARCA4 cell models (doxycycline, 1 µg/mL; 72 h) with or without SAHA treatment for 72 h. Bars show mean ± SD. Two-sided unpaired Student’s t-test. **c**, Western blots depicting levels of indicated proteins in the indicated cells infected with the shNT and with different shSMARCA4 (#1, #4) (see Ref. 7 for more information).

**Extended Data Fig. 3. a**, Heatmaps of normalised ChIP-seq intensities, centred ± 2 kb around the transcriptional start sites (TSS), of SMARCA4, H3K27ac, H3K27me3 and EZH2, in the H1299 cell model. **b**, Genome-wide functional annotations for peaks generated by ChIP-seq analyses. Promoters are defined as the regions ± 2 kb around the annotated TSS. Numbers below each bar indicate the absolute number of peaks at promoters called for the ChIP-seq of each indicated protein and condition. H1299-wtSMARCA4 or H1299-mutSMARCA4 cell models after induction of SMARCA4 (doxycycline, 1 µg/mL; 72 h) with or without SAHA treatment (1 µM, 72 h).

**Extended Data Fig. 4. a**, Venn diagrams of the overlap of SMARCA4, H3K27ac and EZH2 peaks in the H1299 cell model. **b**, Representative snapshots from IGV of ChIP-seq profiles at selected target loci, performed in the H1299 cell model. H1299-wtSMARCA4 or H1299-mutSMARCA4 cell models after induction of SMARCA4 (doxycycline, 1 µg/mL; 72 h) with or without SAHA treatment (1 µM, 72 h).

**Extended Data Fig. 5. a**, Significant correlation (linear regression) between mRNA levels of *KDM6B* (RNA-sequencing from the CCLE database, or RT-QPCR data from current study) and EC_50_ for SAHA (from www.cancerrxgene.org database, or our own data) for each indicated cancer cell line. **b**, Western blots depicting levels of the indicated proteins in the indicated cells infected with the shNT and with different shKDM6 (#60, #61, #62, #63 and #64) and shKDM6B (#676, #976, #75 and #77). **c**, Clonogenic assays for the H460 cells infected with shNT, shKDM6 (#63, #64) or shKDM6B (#676, #976) treated with SAHA for 5 days.

**Extended Data Fig. 6. a**, Examples of the effects on cell viability, measured using MTT assays, after treatment with increasing concentrations of GSK-J4 in the indicated lung cancer cell lines. Lines, percentage of viable cells relative to the total number at 0 h. Error bars, means ± SD of three replicates. **b-c**, clonogenic (**b**) and cell viability (MTT) (**c**) assays in the H1299-wtSMARCA4 or H1299-mutSMARCA4 cell models before and after induction of SMARCA4 (doxycycline, 1 µg/mL; 5 days) treated with GSK-J4. Lines, percentage of viable cells relative to the total number at 0 h. Error bars, mean ± SD of three replicates.

**Extended Data Fig. 7**. Representative sections of H&E staining of the indicated tumours developed from the indicated lung cancer cells grown in nude mice (lung orthotopic implantation of small solid fragments of engrafted cell lines previously grown subcutaneously), treated with GSK-J4 or vehicle. Left panels, scale bars, 2.5 mm. Magnification as indicated (5x, 10x and 20x panels; scale bars, 100 µm, 50 µm and 25 µm, respectively).

**Extended Data Fig. 8. *In vivo* establishment of paired cisplatin-resistant SCCOHT xenografted tumour and effect of cisplatin and GSK-J4 treatments. a**, Western blot to show presence or absence of SMARCA4 in the indicated ovarian cancer cells. **b**, Lateral laparotomy was conducted in isofluorane-anaesthetised mice. The ovary was mobilised and small tumour pieces of primary tumour were anchored on the mouse ovarian surface with prolene 7.0 sutures. Engrafted OVA250X tumours grew as large solid masses, usually to 1,000–1,500 mm^3^ in diameter, which determined the time of sacrifice. The experimental approach used for cisplatin (CDDP)-resistant tumour generation combines: (i) iterative cycles of cisplatin treatment (3 doses of cisplatin administered by i.v. injection on days 0, 7 and 15); (ii) successive increase of administered doses through four cycles applied to different mice. **c**, Representative H&E staining reveals a similar morphology between primary SCCOHT and paired engrafted OVA250X. Top panels, scale bar 500 µm, middle panels scale bar, 200 µm. Bottom panels scale bar, 50 µm. **d**, Gross pathological photographs at necropsy (top panels) and representative H&E staining (bottom panels) of the OVA250X and OVA250XR tumours. Scale bar, 50 µm. **e**, Measure of the weight of each of the tumours from mice treated with vehicle (VH) or cisplatin (CDDP). **f**, Left panel, gross pathological photographs of all ovarian tumours that developed in the mice treated with vehicle (VH), cisplatin (CDP) or GSK-J4. Right panels, volume and weight of each tumour. **g**, Representative H&E staining of OVA250XR tumours, treated as indicated. Scale bar, 100 µm (upper panels) and 25 µm (lower panels). **h**, Quantification of necrotic areas.

All tests are two-sided unpaired Student’s t-test. *, *P* < 0.05; n.s, not significant.

## Methods

### Cancer cell cultures

Cell lines were obtained from the American Type Culture Collection (ATCC), grown under recommended conditions and maintained at 37°C in a humidified atmosphere of 5% CO_2_/95% air. The cells lines were authenticated by genotyping for *TP53* and other known mutations. All cell lines used in this study were mycoplasma-free. Genomic DNA and total RNA were extracted by standard protocols. Two primary cancer cell lines cultures were derived from orthoxenografts/PDOXs generated in nude mice from two primary tumours of two SCCOHT patients (OVA250L and OVA259L) that were obtained from Bellvitge Hospital and the Catalan Institute of Oncology (ICO) with the approval of the Ethical Committee (CEIC Bellvitge). Ethical and legal protection guidelines of human subjects, including informed consent, were followed. Fresh orthoxenografts/PDOXs grown in the mouse ovaries were collected when mice were sacrificed at passage #1, then minced with sterile scalpels. Single cells and clumps were transferred to cell culture plates and maintained in DMEM supplemented with 10% FBS plus 50 U/mL penicillin and 50 mg/mL streptomycin under standard culture conditions. When cell colonies with epithelial cell morphology were observed, cells were trypsinized and expanded. Both primary cell lines were considered established after > 6 passages *in vitro*. Specific informed consent was obtained from all patients for tumour implantation into mice, and the study was approved by the IDIBELL Ethics Committee (No. AAALAC-1155).

### Antibodies and western blots

The following primary antibodies were used for western blots: anti-TUBULIN, T6199 mouse (1/10000, Sigma Aldrich, St. Louis, MO, USA); anti-Beta-ACTIN, 13854 (1/20000 Sigma Aldrich); anti-SMARCA4 49360S (1:1000, Cell Signaling Technology); anti-EZH2 5246S (1:1000, Cell Signaling); anti-H3K27ac D5E4 (1:1000, Cell Signaling) anti-H3K27me3 07-449 (1:1000, Cell Signaling); anti-UTX (KDM6A) D3Q1I (1:1000, Cell Signaling); anti-KDM6B (JMJD3) AB154126 (1:1000 Abcam) (see also Extended Supplementary Tables). For western blots, whole-cell lysates were collected in a buffer containing 2% SDS 50 mM Tris-HCl (pH 7.4), 10% glycerol and protease inhibitor cocktail (Roche Applied Science). Protein concentrations were determined using a Bio-Rad DC Protein Assay kit (Life Science Research). Equal amounts of lysates (20 µg) were separated by SDS-PAGE and transferred to a nitrocellulose membrane that was blocked with 5% nonfat dry milk. Membranes were incubated with the primary antibody overnight at 4°C, then washed before incubation with species-appropriate IRDye 680CW (925-68022) or IRDye 800CW (925-32213) fluorescent secondary antibodies (1:10.000 LI-COR, NE, USA) for 1 h at room temperature.

### Treatments and shRNAs

Chemicals were obtained from the following sources: SAHA, suberoyl anilide hydroxamic acid (Cayman Chemical Company, Ann Arbour, MI, USA); GSK-J4 (Shelleckchem); GSK-126 (Cayman Chemical Company). shRNAs against SMARCA4, KDM6a and KDM6B were purchased from SIGMA-MISSION (LentiExpressTM Technology, Sigma-Aldrich) as a glycerol stock of five pLKO plasmids carrying specific shRNA sequences. A non-target shRNA (shNT) (Sigma MISSION shRNA non-mammalian control SHC002) was used as a control. The lentiviruses were generated within the 293T packaging cells.

### Cell growth analysis and calculation of EC_50_ and combination index (CI)

For cell viability assays, cell lines were incubated in 96-well plates. Before harvesting, cells were treated for 5-7 days with the indicated concentrations of each compound (SAHA, GSK-J4, GSK-126) or combinations. For the assays, 10 µl of a solution of 5 mg/mL MTT [3-(4,5)-dimethylthiazol-2-yl)-2,5-diphenyltetrazolium bromide] (Sigma Chemical Co.) was added. After incubation for 3 h at 37°C, the medium was discarded, the formazan crystals that had formed were dissolved in 100 µl of lysis buffer (50% N-N-dimethylformamide in H_2_O, 20% SDS, 2.5% glacial acetic acid, NaOH 5 mol/L, pH 4.7), and absorbance was measured at 596 nm. Results are presented as the median of at least two independent experiments performed in triplicate for each cell line and for each condition. For EC_50_ calculations, cells were treated with each drug and their various combinations for 5 days. Estimates of EC_50_ were derived from the dose response curves. To assess the drug concentration effect and to calculate the combination index (CI), cells were plated in 96-well plates and incubated with a concentration of SAHA ranging from 0.07 μM to 10 μM (0.07, 0.15, 0.3, 0.6, 1.25, 2.5, 5.0 and 10 μM), and the same for GSK-J4 for 5 days. MTT assays were performed and the EC_50_ was determined by fitting the dose-response curve utilizing the CompuSyn software. The CI values for each dose and the corresponding effect level were calculated. The combination index offers a quantitative definition for drug combinations in which CI < 1, CI = 1 and CI > 1 indicate synergism, an additive effect, and antagonism, respectively.

For clonogenic assays, the plates were seeded with 5,000 cells from each cell line, then treated with SAHA (1 µM), GSK-J4 (1 µM) or GSK-126 (1 µM) for 5 days. Cells were stained with crystal violet solution (0.5% Crystal Violet in 25% of methanol).

### Quantitative RT-PCR

To assess mRNA levels of the KDMs in different cells qPCR analysis was performed. 1 μg of total RNA was reverse-transcribed using SuperScript™ II reverse transcriptase (Invitrogen) and Random primers (Promega), according to the manufacturers’ instructions. qRT-PCR was performed in a Quantstudio 5 Real-Time PCR Instrument using SYBR Green PCR Master Mix (Applied Biosystems). Three biological replicates were carried out. Primer sequences are provided in the Extended Supplementary Tables.

### ChIP-sequencing

For ChIP, cells were grown in P-150 cm cell dishes and fixed with 1% methanol-free formaldehyde (Thermo Scientific) for 10 min at room temperature, then quenched by 125 mmol/L glycine for 15 min at room temperature, washed with ice-cold PBS twice and centrifuged at 200g, 4°C for 5 min. The pellet was resuspended in 1 mL of cell lysis buffer (10 mmol/L Tris-HCl, 10 mmol/L NaCl, 0.5% NP-40, protease inhibitor) and kept at 4°C, rotating for 30 min. After centrifugation, the pellet was resuspended in 1 mL of nuclear lysis buffer (1% SDS, 10 mmol/L EDTA, 50 mmol/L Tris-HCl pH 8.0, protease inhibitor) and kept at 4°C for 60 min. After another centrifugation, the lysate was sonicated with a Covaris M220 instrument to yield chromatin fragments of an average size of 0.25–1.00 kb, and then frozen at -20°C for 30 min. The chromatin was thawed on ice and centrifuged at 2,500 g. For each ChIP reaction, 60 μL of Magna ChIP™ Protein A+G Magnetic Beads (Merck Millipore) was used in accordance with the manufacturer’s protocol. Before addition of the sheared chromatin to the beads, Triton X-100 and Na-deoxycholate was added to a final concentration of 10% each. 1% of the chromatin volume was used for input. At least two independent ChIP experiments were performed.

Immunoprecipitated chromatin was deep-sequenced in the Genomics Unit of the Centre for Genomic Regulation (CRG, Barcelona, Spain) using the Illumina HiSeq 2500 system (Illumina). Briefly, library preparation included end-repair, generation of dA overhangs, adapter ligation, size selection and removal of non-ligated adapters by agarose gene electrophoresis and amplification (18 cycles) before loading the samples into the sequencer.

For ChIP-sequencing data analysis, reads were aligned to the human reference genome hg38, using Bowtie v1.2.2, with default parameters and disallowing multi-mapping (–m 1)^35^. PCR duplicates were removed using PICARD (http://broadinstitute.github.io/picard/). Ambiguous and multi-mapped reads were discarded. Peaks were called using MACS2 v2.1.1^36^. To avoid false positives, peaks were discarded if they were present in the ChIP-seq of SMARCA4 in the SMARCA4-deficient cells. Genomic peak annotation was performed with the R package ChIPpeakAnno v3.15, considering the region ± 2 kb around the TSS as the promoter^37^. All analyses considered peaks overlapping with promoter regions, unless otherwise specified. Peak lists were then transformed to gene target lists.

For heatmap and intensity plot representation of ChIP-seq signal, bedgraph files were generated using the makeUCSCfile function in HOMER with default parameters normalizing for differences in sample library size, and BigWig files were generated using the function bedGraphToBigWig from UCSC. Heatmaps were derived using the functions computeMatrix, in a window of ± 2 kb around the centre in the TSS, and plotHeatmap from deepTools^38^.

### Generation of orthotopic tumour models and treatments

Male and female athymic nu/nu mice (ENVIGO) aged 4-5 weeks were maintained in a sterile environment before use in the lung cancer orthotopic experiments. Female athymic nu/nu mice (ENVIGO), 4-6 weeks old, were used for the ovarian cancer orthotopic studies. The animals were housed in individually ventilated cages on a 12-hour light-dark cycle at 21-23°C and 40-60% humidity. Mice were allowed free access to an irradiated diet and sterilized water. All animal experiments were approved by the IDIBELL Ethical Committee under protocol 9111 approved by the Government of Catalonian, AAALAC accredited Unit 1155, and performed in accordance with guidelines stated in the International Guiding Principles for Biomedical Research Involving Animals, developed by the Council for International Organizations of Medical Sciences (CIOMS). To generate orthotopic lung tumour xenografts the cell lines were injected subcutaneously into the back of the mice (n = 3 mice/cell line). Once the solid tumour had entered the exponential growth phase, mice were sacrificed, the tumour was isolated under sterile conditions, and the non-necrotic areas were selected and minced in small fragments of 2-3 mm^3^. These were then orthotopically implanted in the lung parenchyma, as previously described^20,27^. On day 15, the mice were randomized and intraperitoneally treated with GSK-J4 (50 mg/kg/day for each mouse) or corresponding vehicle only. For the lung orthotopic models, in most cases the animals were sacrificed when they displayed serious respiratory difficulty, which was subsequently confirmed to be associated with lung tumour growth

Orthoxenografts or patient-derived orthotopic xenografts of small cell carcinoma of the ovary, hypercalcaemic type (SCCOHT) were generated. The primary tumour specimens for the two primary SCCOHT samples were freshly obtained at Hospital Universitari de Bellvitge (Hospitalet de Llobregat, Barcelona, Spain). The study was approved by the Institutional Review Board, and written informed consent was obtained from both patients. The orthotopic ovarian tumours were engrafted in mice, following an established protocol^27^. Briefly, non-necrotic tissue pieces (2–3 mm^3^) from resected carcinoma were selected and placed in DMEM (BioWhittaker) supplemented with 10% FBS and penicillin/streptomycin at room temperature. Under isofluorane-induced anaesthesia, animals were subjected to a lateral laparotomy, their ovaries exposed and tumour pieces anchored to the ovary surface with prolene 7.0 sutures. Tumour growth was monitored 2 or 3 times per week and when the tumour had reached a sufficient size, it was harvested, cut into small fragments, and transplanted into between two and four new animals. Engrafted tumours (named OVA250X) at early mouse passages were cut into 6-8 mm^3^ pieces and stored in liquid nitrogen in a cryopreservation solution of 90% FBS and 10% dimethyl sulfoxide, awaiting subsequent implantation.

To generate the cisplatin-resistant ovarian xenograft mouse model, orthotopically engrafted OVA250X tumours at passage#1 were allowed to grow (n=3 mice) until palpable intra-abdominal masses were noted. Cisplatin was i.v.-administered to the animals (cycle #1, 3, 5 mg/kg dose) for 3 consecutive weeks (days 0, 7 and 15; cycle#1 of treatment) (Extended Data Fig. 8b). Post-cisplatin tumours at relapse were harvested and engrafted in new animals. This process of cisplatin treatment was repeated up to four times by treating tumour-bearing mice with stepwise-incremental doses of cisplatin: cycle #2, 4 mg/kg; cycle #3, 5 mg/kg and cycle #4, 5 mg/kg (see Extended Data Fig. 8b). Cisplatin-resistant tumours were obtained (OVA250XR). At doses higher than 3.5 mg/kg, signs of cisplatin induced some toxicity that were ameliorated by 2 days administration of saline containing 5% glucose. Mice were transplanted with fragments of OVA250X and OVA250XR tumours, and when tumours reached a homogeneous palpable size were randomly allocated into the treatment groups (n = 3-7 mice/group): i) Placebo; ii) GSK-J4 (50 mg/kg) and iii) cisplatin (3.5 mg/kg); drugs were administered once a day, 5 days per week, for 4 consecutive weeks. Animals were sacrificed on day 21 of treatment, and their ovaries dissected out and weighed. Representative fragments were either frozen in nitrogen or fixed and processed for paraffin embedding.

### Histopathology and immunostaining

For histological analysis, tumours were fixed and embedded in paraffin. Necrosis/fibrosis were morphological assessed after staining with haematoxylin and eosin (H&E), using standard protocols, and then examined by light microscopy in a blinded fashion. For immunostainings, 4-μm-thick paraffin-embedded sections of lung and ovarian tumour samples were deparaffinized overnight at 62°C and then immersed in xylene. Samples were rehydrated and, after microwaving with Tris/EDTA pH 9.0 for antigen retrieval, endogenous peroxidase was inhibited with a 3% hydrogen peroxide solution, blocked in 10% goat serum and incubated with primary antibodies overnight at 4°C (Extended Supplementary Tables). HRP-conjugated polyclonal goat (anti-mouse and anti-rabbit) secondary antibodies (NeoStain ABC Kit, NeoBiotech) were used in 1-h incubations at room temperature. Labelling detection was done using an ImmPACT DAB Peroxidase (HRP) Substrate kit (Vector Laboratories, Burlingame, CA, USA), and tissue sections were counterstained with haematoxylin. Once dehydrated in an ethanol battery and cleared in xylene for 1 h, samples were mounted with coverslips with DPX mounting medium (Merck Millipore, Darmstadt, Germany). Sections were evaluated under a Leica DM1000 microscope by two independent observers in a blind fashion. Areas of necrosis/fibrosis were quantified using Photoshop. The scoring criteria for determining H3K27me3 staining were based on the staining intensity (four categories, 1-4). The mean of values from three independent evaluators was determined.

### Statistical analysis

Student’s t-tests, EC_50_ calculations, Kaplan-Meier estimates and log-rank (Mantel-Cox) test were performed using Prism software (GraphPad). Values of *P* less than 5% were considered statistically significant. The statistical methods used for each analysis are specified in the figure legends.

## Data availability

The ChIP-seq data obtained in this study have been uploaded to the SRA (NCBI), under accession number (Pending).

## Acknowledgments

The authors thank Isabel Bartolessis (Cancer Genetics Group) at IJC for technical assistance. This work was supported by the Spanish Ministry of Economy and Competitivity-MINECO (grant number SAF-2017-82186R, to M. Sanchez-Cespedes, and grant PI19/01293 to A. Villanueva) and from the Fundación Científica of the Asociación Española Contra el Cancer (AECC) (grant number GCB14142170MONT) to M. Sanchez-Cespedes. A. Villanueva is also funded by the Department of Health of the Generalitat de Catalunya (2014SGR364). O.A. Romero received a Juan de la Cierva postdoctoral contract (grant No. IJCI-2016-28201, until November 2019) and an AECC research contract (INVES19045ROME from December 2019). A. Vilarrubi, P. Llabata and A. Andrades are supported by pre-doctoral contracts from the Spanish MINECO (FPI-fellowship: PRE2018-084624, BES-2015-072204 and FPU17/00067). M. Saigi was supported by a Rio Hortega contract from the Instituto de Salud Carlos III (CM17/00180). L. Farre received a European Union Horizon 2020 research and innovation programme under the Marie Sklodowska-Curie Actions grant agreement, number 799850.

## References

1. Clapier, C.R. & Cairns, B.R. The biology of chromatin remodeling complexes. Annu. Rev. Biochem. 78, 273–304 (2009).

2. Mohrmann, L. & Verrijzer, C.P. Composition and functional speci?city ofSWI2/SNF2 class chromatin remodeling complexes. Biochim. Biophys. Acta 1681, 59–73 (2005).

3. Romero, O.A. & Sanchez-Cespedes, M. The SWI/SNF genetic blockade: effects in cell differentiation, cancer and developmental diseases. Oncogene 33, 2681–2689(2014).

4. -Medina, P.P. et al. Frequent BRG1/SMARCA4-inactivating mutations in human lung cancer cell lines. Hum. Mut. 29, 617–622 (2008).

5. Rodriguez-Nieto, S. et al. Massive parallel DNA pyrosequencing analysis of the tumor suppressor BRG1/SMARCA4 in lung primary tumors. Hum. Mut. 32, E1999–2017 (2011).

6. Romero, O.A. et al. The tumour suppressor and chromatin-remodelling factor BRG1 antagonizes Myc activity and promotes cell differentiation in human cancer. EMBO Mol. Med.4, 603–616 (2012).

7. -Romero, O.A. et al. MAX inactivation in small cell lung cancer disrupts MYC-SWI/SNF programs and is synthetic lethal with BRG1. Cancer Discov. 4, 292–303 (2014).

8. Matias-Guiu, X, et al. Human parathyroid hormone related protein in ovarian small cell carcinoma. An immunohistochemical study. Cancer 73, 1878–1881 (1994).

9. -Jelinic, P. et al. Recurrent SMARCA4 mutations in small cell carcinoma of the ovary. Nat. Genet. 46, 424–426 (2014).

10. -Witkowski, L. et al. Germline and somatic SMARCA4 mutations characterize small cell carcinoma of the ovary, hypercalcemic type. Nat. Genet. 46, 438–443 (2014).

11. Ramos, P. et al. Small cell carcinoma of the ovary, hypercalcemic type, displays frequent inactivating germline and somatic mutations in SMARCA4. Nat. Genet. 46, 427–429 (2014).

12. -Oike, T. et al. A synthetic lethality-based strategy to treat cancers harboring a genetic deficiency in the chromatin remodeling factor BRG1. Cancer Res. 73, 5508–5518 (2013).

13. Hoffman, G.R., et al. Functional epigenetics approach identifies BRM/SMARCA2 as a critical synthetic lethal target in BRG1-deficient cancers. Proc. Natl. Acad. Sci. USA. 111, 3128–3133 (2014).

14. Kim, K.H. et al. SWI/SNF-mutant cancers depend on catalytic and non-catalytic activity of EZH2. Nat. Med. 21, 1491–1496 (2015).

15. Xue, Y. et al. SMARCA4 loss is synthetic lethal with CDK4/6 inhibition in non-small cell lung cancer. Nat. Commun. 10, 557 (2019).

16. Chiba, H., Muramatsu, M., Nomoto, A. & Kato, H. Two human homologues of Saccharomyces cerevisiae SWI2/SNF2 and Drosophila brahma are transcriptional coactivators cooperating with the estrogen receptor and the retinoic acid receptor. Nucleic Acids Res. 22, 1815–1820 (1994).

17. De la Serna, I.L., Carlson, K.A. & Imbalzano, A.N. Mammalian SWI/SNF complexes promote MyoD-mediated muscle differentiation. Nat. Genet. 27, 187–190 (2001).

18. -Chi, T.H. et al. Sequential roles of Brg1, the ATPase subunit of BAF chromatin remodeling complexes, in thymocyte development. Immunity 19, 169–182 (2003).

19. -Seo, S., Richardson, G.A. & Kroll, K.L. The SWI/SNF chromatin remodeling protein Brg1 is required for vertebrate neurogenesis and mediates transactivation of Ngn and NeuroD. Development 132, 105–151 (2005).

20. -Romero, O.A. et al. Sensitization of retinoids and corticoids to epigenetic drugs in MYC-activated lung cancers by antitumor reprogramming. Oncogene 36, 1287–1296 (2017).

21. -Richon, V.M., Sandhoff, T.W., Rifkind, R.A. & Marks, P.A. Histone deacetylase inhibitor selectively induces p21WAF1 expression and gene-associated histone acetylation. Proc. Natl. Acad. Sci. USA. 97, 10014–10019 (2000).

22. Tessarz, P. & Kouzarides, T. Histone core modifications regulating nucleosome structure and dynamics. Nat. Rev. Mol. Cell Biol. 15, 703–708 (2014).

23. -Black, J.C., Van Rechem, C. & Whetstine, J.R. Histone lysine methylation dynamics: establishment, regulation, and biological impact. Mol. Cell. 48, 491–507 (2012).

24. Hodges, H.C. et al. Smarca4 ATPase mutations disrupt direct eviction of PRC1 from chromatin. Nat. Genet. 49, 282–288 (2017).

25. Chandrasekaran, R. & Thompson, M. Polybromo-1-bromodomains bind histone H3 at specific acetyl-lysine positions Biochem. Biophys. Res. Commun. 355, 661–665 (2007).

26. -Kruidenier, L. et al. A selective jumonji H3K27 demethylase inhibitor modulates the proinflammatory macrophage response. Nature 488, 404–408 (2012).

27. -Ambrogio, C., et al. Combined inhibition of DDR1 and Notch signaling is a therapeutic strategy for KRAS-driven lung adenocarcinoma. Nat. Med. 22, 270–277 (2016).

28. -Vidal, A. et al. Lurbinectedin (PM01183), a new DNA minor groove binder, inhibits growth of orthotopic primary graft of cisplatin-resistant epithelial ovarian cancer. Clin. Cancer Res. 18, 5399–5411 (2012).

29. Li, Y. & Seto, E. HDACs and HDAC inhibitors in cancer development and therapy. Cold Spring Harb. Perspect. Med. 6, a026831 (2016).

30. -Arcipowski, K.M., Martinez, C.A. & Ntziachristos, P. Histone demethylases in physiology and cancer: a tale of two enzymes, JMJD3 and UTX. Curr. Opin. Genet. Dev. 36, 59–67(2016).

31. Bogershausen, N. & Wollnik, B. Unmasking Kabuki syndrome. Clin Genet. 83, 201–211 (2013).

32. -Gozdecka, M., et al. UTX-mediated enhancer and chromatin remodeling suppresses myeloid leukemogenesis through noncatalytic inverse regulation of ETS and GATA programs. Nat. Genet. 50, 883–894 (2018).

33. -Ntziachristos, P.et al. Contrasting roles of histone 3 lysine 27 demethylases in acute lymphoblastic leukaemia. Nature 514, 513–517 (2014).

34. Hashizume, R. et al. Pharmacologic inhibition of histone demethylation as a therapy for pediatric brainstem glioma. Nat. Med. 20, 1394–1396 (2014).

## References

35. -Langmead, B., Trapnell, C., Pop, M. & Salzberg, S.L. Ultrafast and memory-efficient alignment of short DNA sequences to the human genome. Genome Biol. 10, R25 (2009).

36. -Zhang, Y. et al. Model-based Analysis of ChIP-Seq (MACS). Genome Biol. 9, R137 (2008).

37. - Zhu, L. et al. ChIPpeakAnno: a Bioconductor package to annotate ChIP-seq and ChIP-chip data. BMC Bioinformatics 11, 237 (2010).

38. -Ramírez, F. et al. deepTools2: A next generation web server for deep-sequencing data analysis. Nucleic Acids Res. 44, w160-165 (2016).

