## Supplementary Tables for "SMARCA4 deficient tumours are vulnerable to KDM6A/UTX and KDM6B/JMJD3 blockade"

**Extended data Table 1**. List of antibodies, including source, concentrations and analysis. WB, western-blot; IHC, immunohistochemistry; ChIP, chrommatin immunoprecipitation.

| Protein | Reference | Supplier | Technique | Dilution (WB) | Total amount (ChIP) | Dilution (IHC) |
|---|---|---|---|---|---|---|
| KDM6B/JMJD3 | ab154126 | Abcam | WB | 1:1000 | | |
| | ab38113 | Abcam | WB & IHC | 1:1000 | | 1:50 |
| KDM6A/UTX | D3Q1I | Cell signaling | WB & IHC | 1:1000 | | 1:100 |
| H3K27me3 | 07-449 | Cell signaling | WB & ChIP-seq & IHC | 1:1000 | 2 micrograms | 1:100 |
| H3K27ac | D5E4 | Cell signaling | WB & ChIP-seq & IHC | 1:1000 | 2 micrograms | 1:100 |
| EZH2 | 5246S | Cell signaling | WB & ChIP-seq | 1:1000 | 2 micrograms | |
| SMARCA4 | 49360S | Cell signaling | WB & ChIP-seq | 1:1000 | 2 micrograms | |
| ACTIN | 13859 | Sigma | WB | 1:10000 | | |
| TUBULIN | T6199 | Sigma | WB | 1:10000 | | |

**Extended data Table 2**. List of primers used for the quantitative RT-PCRs.

| Gene | Forward (5'-3') | Reverse (5'-3') | Scale (µmoles) | Supplier |
|---|---|---|---|---|
| *ACTB* | GGCATCCTCACCCTGAGGTA | AGGTGTGGTCGGAGATTTTC | 0.025 | Sigma |
| *GUSB* | CTGTACACGACACCCACCAC | GATGAGGAACTGGCTCTTGA | 0.025 | Sigma |
| *KDM1A/LSD1* | GCCATGGTGGTACAGGTCTA | TGGCCAGTTCCATATTTACA | 0.025 | Sigma |
| *KDM2A* | CGGATAGTTGAGAAAGCCAT | CTCTTTGGTGGGCCTCTGTA | 0.025 | Sigma |
| *KDM3A* | ACCTGCAGTTATTCTTCAGC | TAATGCCAGTCCTATGCCAT | 0.025 | Sigma |
| *KDM4A/JMJD2A* | CCTCACTGCGCTGTCTGTAT | CCAGTCGAAGTGAAGCACAT | 0.025 | Sigma |
| *KDM4B/JMJD2B* | GATCTCCATGGACGTGTTCG | CGTTTCCGGTGAGACCTGCC | 0.025 | Sigma |
| *KDM4C/JMJD2C* | GGCATAGGTGACAGGGTGTG | CGGGGACCAAACTCTGGAAA | 0.025 | Sigma |
| *KDM4D/JMJD2D* | CGGGATCTGCACAGATTATC | AGTTTCTGAGGAGGGCGACC | 0.025 | Sigma |
| *KDM5A* | GGTTTCTTAAGGTGGCAAGT | TTCTTTTGTACTGTTCCCTA | 0.025 | Sigma |
| *KDM5B* | AGCTTTTCTCAGAATGTTGG | GCAGAGTCTGGGAATTCACA | 0.025 | Sigma |
| *KDM5C* | GGGTTTCTAAAGTGTAGATC | CCCACACATCTGAGCTTTAG | 0.025 | Sigma |
| *KDM6A/UTX* | GTCCGAGTGTCAACCAACTG | TGAGAGTCCTGGAGTAGGAG | 0.025 | Sigma |
| *KDM6B/JMJD3* | CTCAACTTGGGCCTCTTCTC | GCCTGTCAGATCCCAGTTCT | 0.025 | Sigma |

**Extended data Table 3**. References for the shRNAs

| Name | Clone ID | Vector | Target gene | Supplier |
|---|---|---|---|---|
| #60 | TRCN0000107760 | pLKO.1 | *KDM6A* | Sigma |
| #61 | TRCN0000107761 | pLKO.1 | *KDM6A* | Sigma |
| #63 | TRCN0000107763 | pLKO.1 | *KDM6A* | Sigma |
| #64 | TRCN0000107764 | pLKO.1 | *KDM6A* | Sigma |
| #77 | TRCN0000236677 | pLKO.1 | *KDM6B* | Sigma |
| #78 | TRCN0000236678 | pLKO.1 | *KDM6B* | Sigma |
| #676 | TRCN0000236676 | pLKO.1 | *KDM6B* | Sigma |
| #976 | TRCN0000359976 | pLKO.1 | *KDM6B* | Sigma |
