## Supplementary Figures for "SMARCA4 deficient tumours are vulnerable to KDM6A/UTX and KDM6B/JMJD3 blockade"

### Extended Data Fig.1

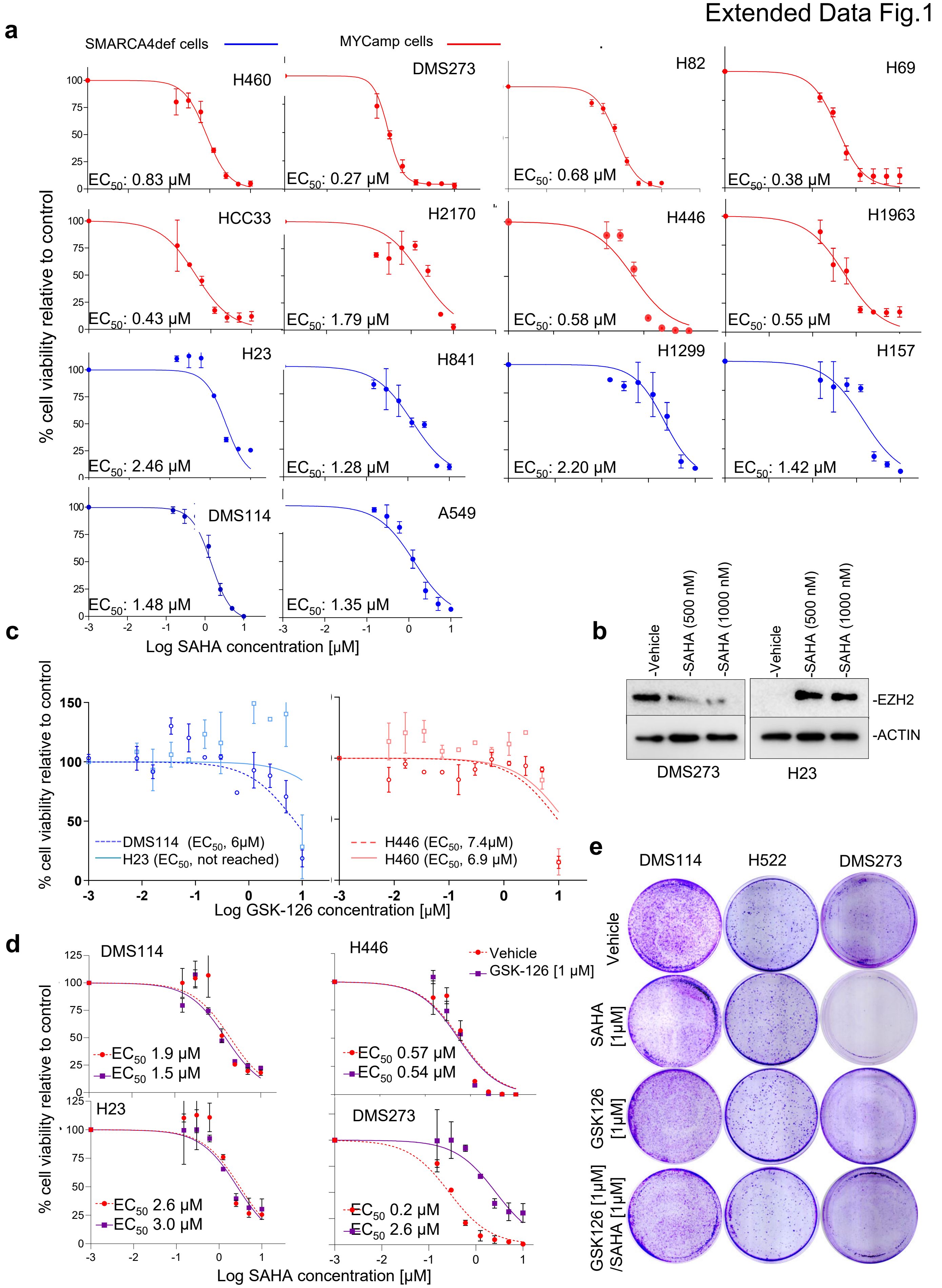

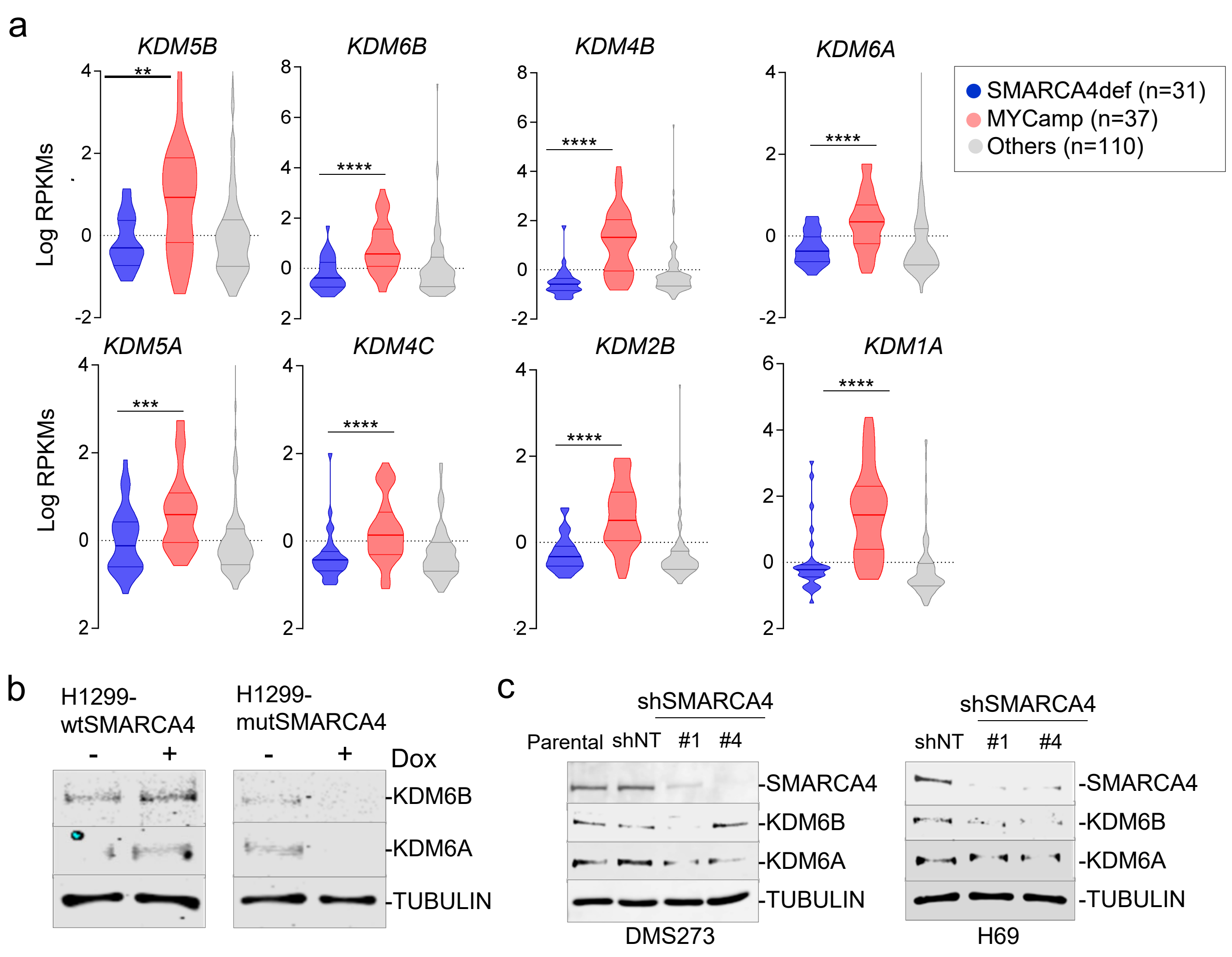


Extended Data Fig.2

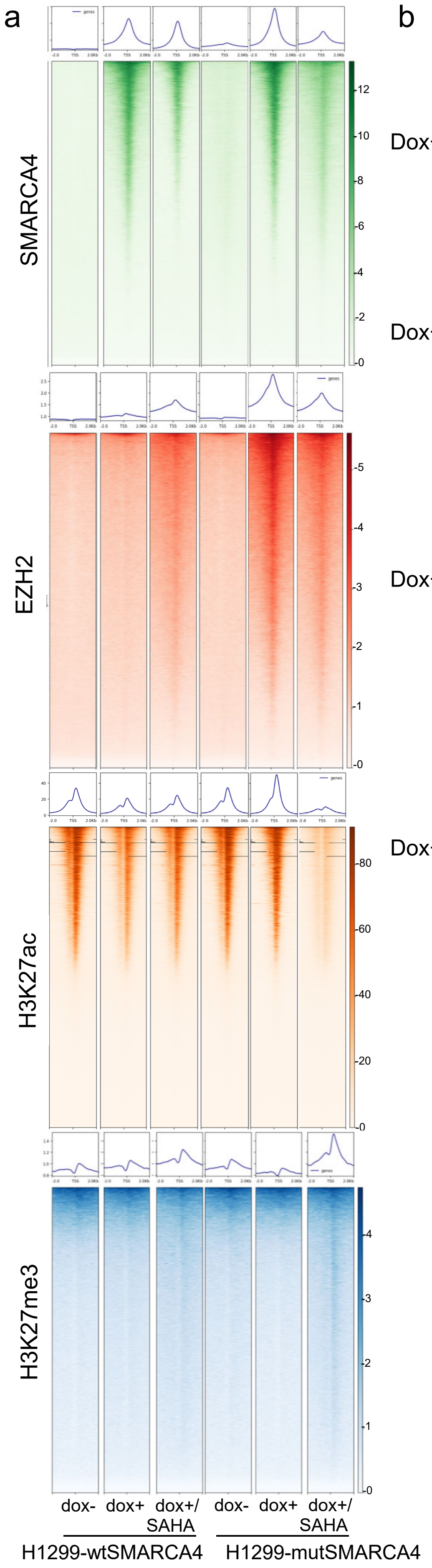


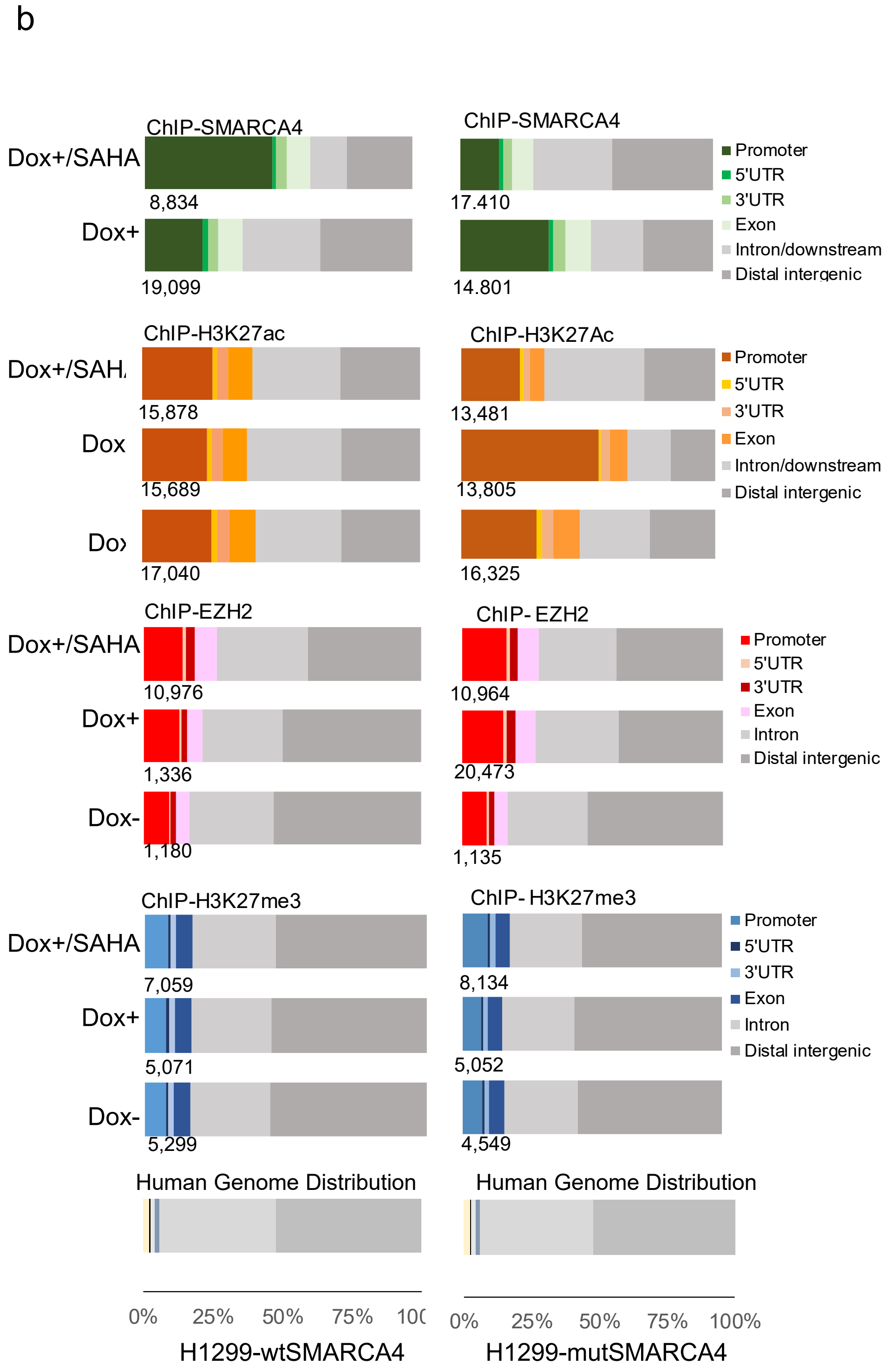


Extended Data Fig.3

### Extended Data Fig.4

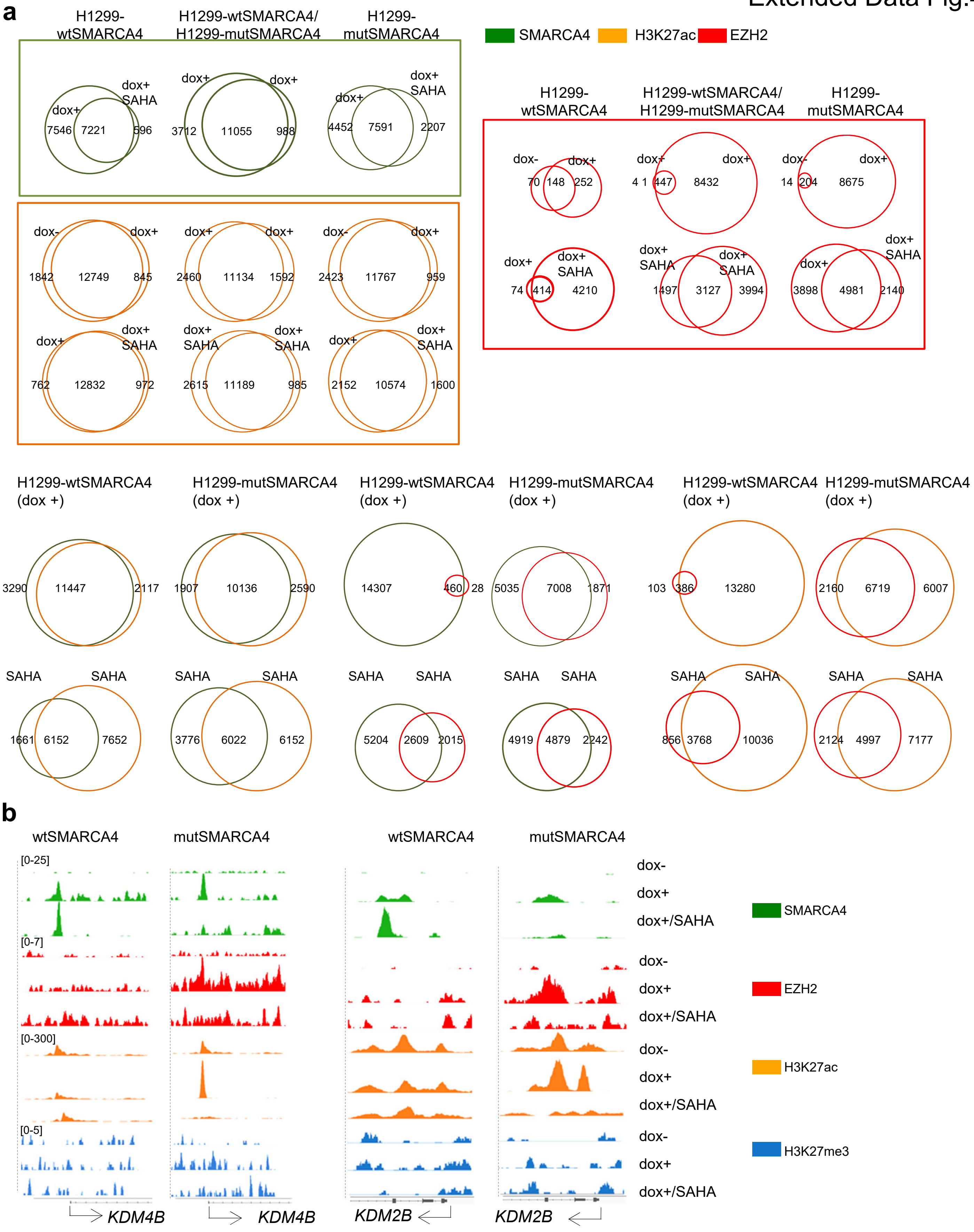

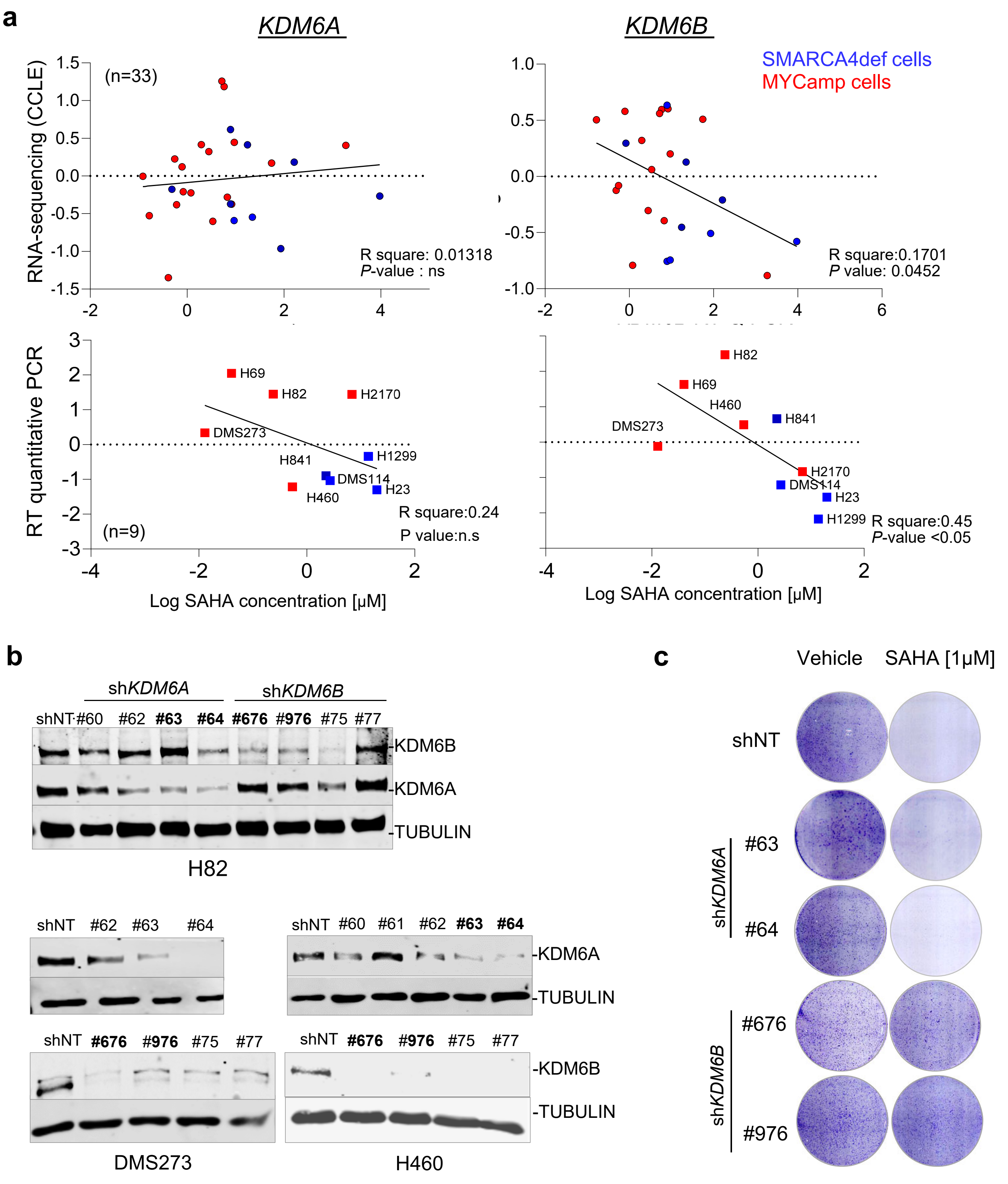


Extended Data Fig.5

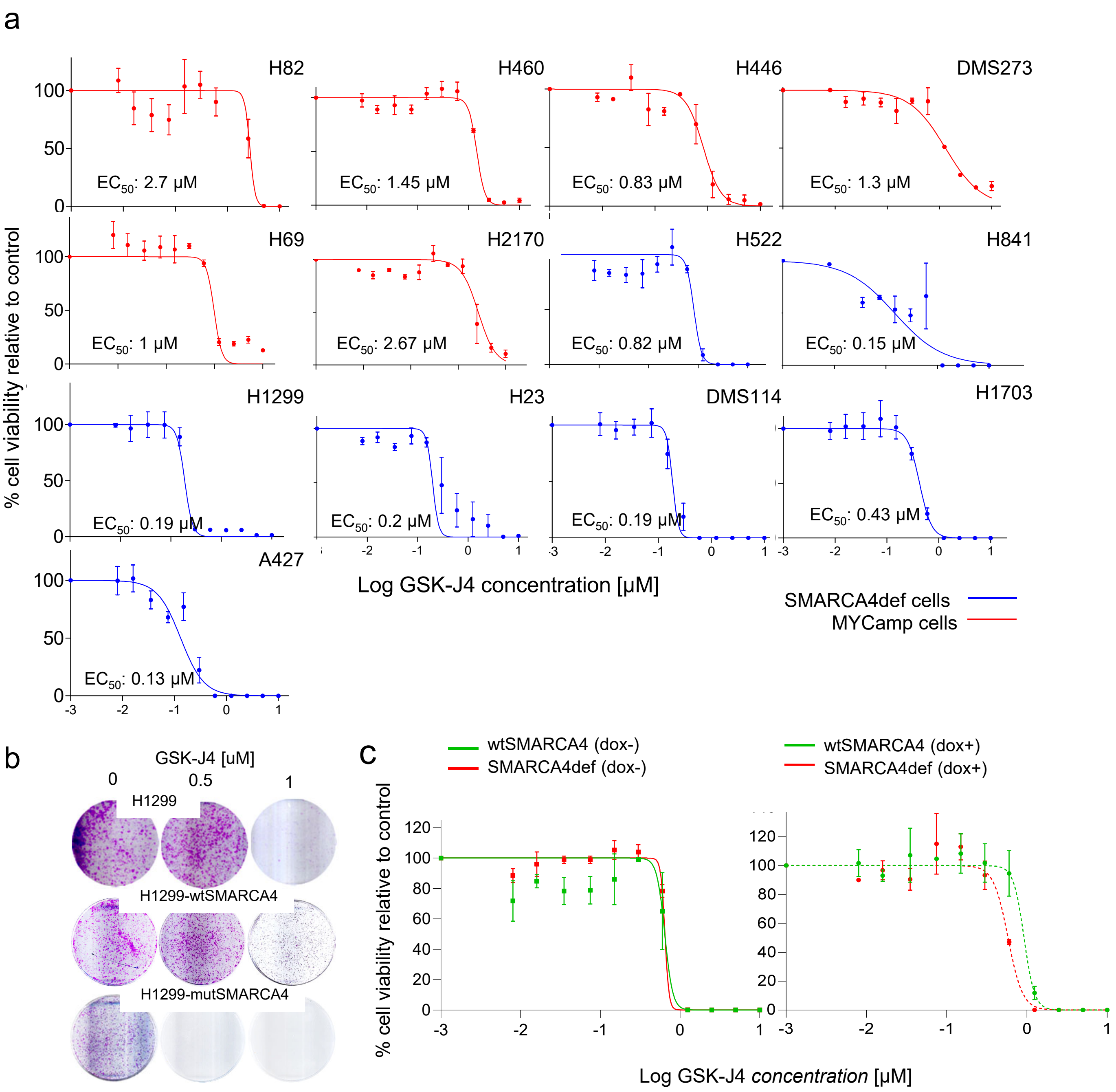


Extended Data Fig.6

Extended Data Fig.7

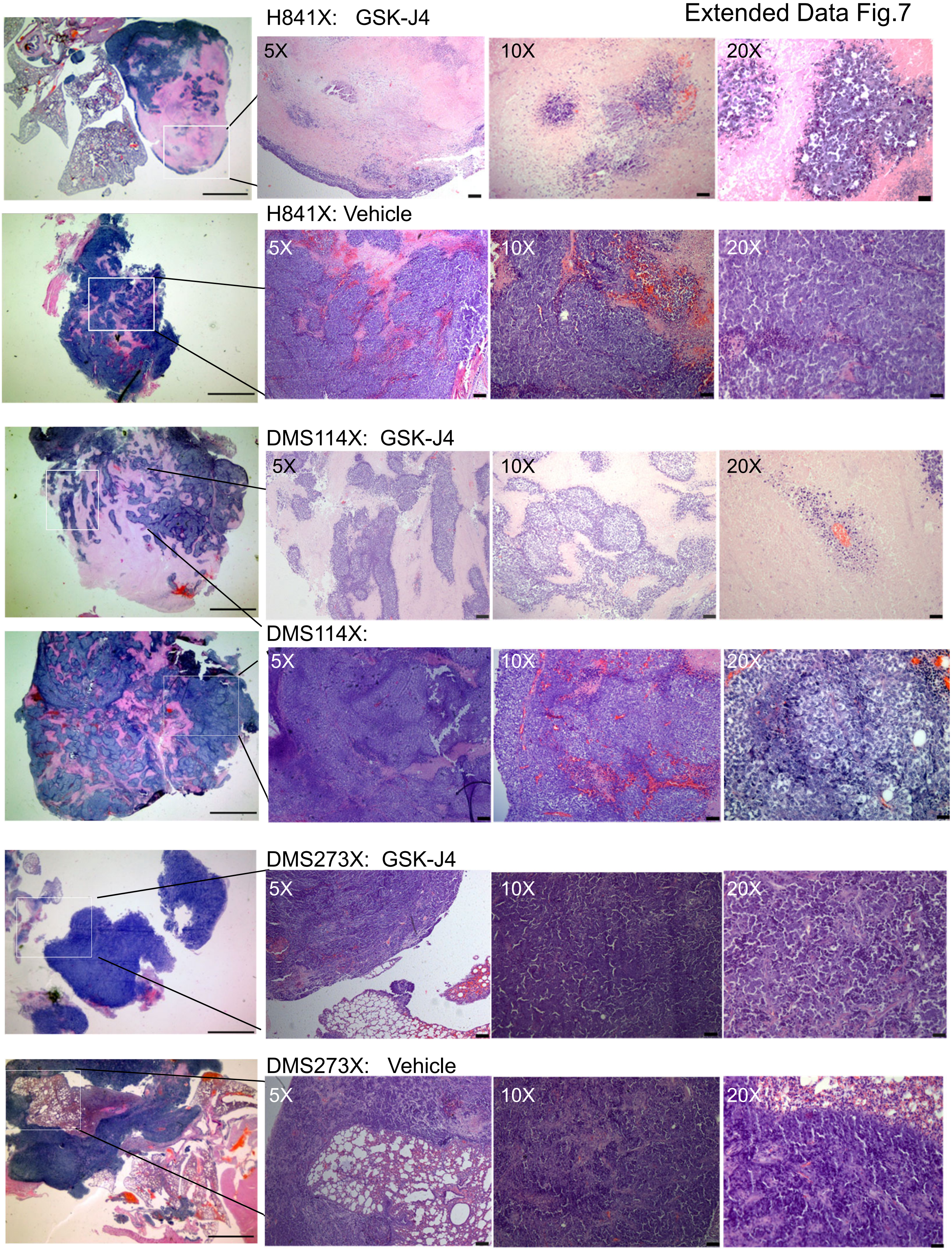

Extended Data Fig.8

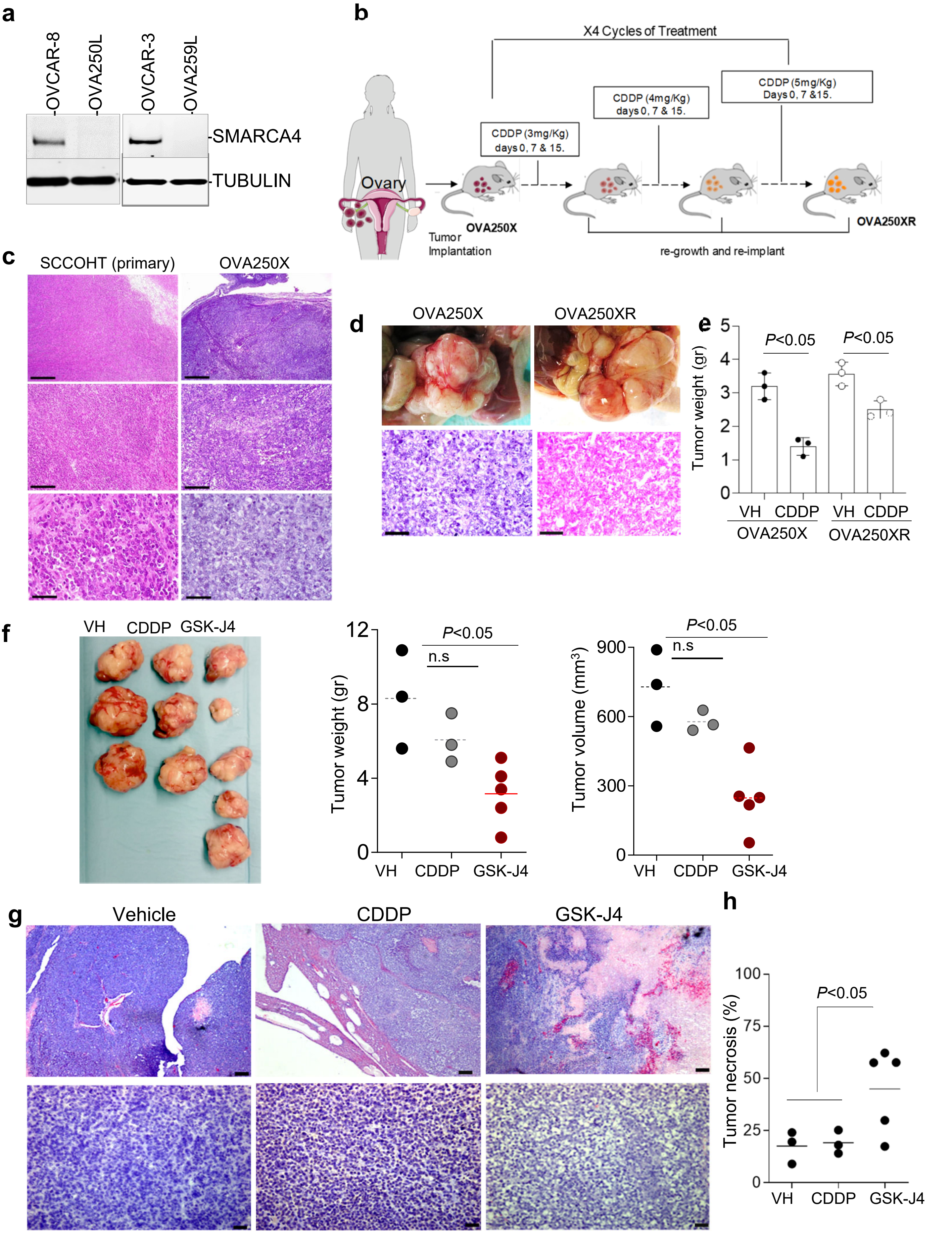
